# Cryo-EM Structures of Apo, Agonist- and Antagonist-Bound Heteromeric Kainate Receptors

**DOI:** 10.64898/2025.12.02.691905

**Authors:** Changping Zhou, Guadalupe Segura-Covarrubias, Ashlee M’Kay Hoffman, Nami Tajima

## Abstract

Kainate receptors (KARs) mediate excitatory synaptic transmission and regulate neurotransmitter release. In the central nervous system, KARs predominantly exist as heterotetramers comprising low-affinity (GluK1–3) and high-affinity (GluK4–5) subunits, with GluK2/GluK5 being the most abundant. To elucidate their conformational transitions, we determined the cryo-electron microscopy (cryo-EM) structures of GluK2/GluK5 KARs in the apo and distinct ligand-bound states. The apo structure revealed compact packing with extensive inter-subunit interactions between the ligand-binding domains (LBDs) beyond the conserved D1-D1 upper-lobe contacts. The glutamate-bound structure exhibited enhanced packing that stabilized the desensitized conformation through increased intersubunit contacts relative to the homomeric KARs, indicating that the heterotetramers were conformationally less dynamic. To investigate subtype-specific antagonism, we engineered GluK2 and GluK5 mutants with altered affinities for the competitive antagonist UBP310. Structural analysis of these mutants revealed distinct UBP310 binding modes on the GluK2 versus the GluK5 subunits. Furthermore, we demonstrated that targeting GluK5 was more effective than targeting GluK2. Specifically, blocking GluK5 locked the receptor in a pore-occluded conformation, whereas antagonizing GluK2 retained structural flexibility above the pore. These findings provide a structural framework for understanding the distinct contributions of the GluK2 and GluK5 subunits to KAR function.

**Teaser:** Cryo-EM structures of heteromeric kainate receptors reveal unique assembly and regulation mechanisms.

## Introduction

Kainate receptors (KARs) are a subfamily of ionotropic glutamate receptors (iGluRs), which are widely expressed throughout the central nervous system and contribute to synaptic structure and function(*1*). They localize pre-, post-, and extrasynaptically and mediate signaling through ionotropic or metabotropic mechanisms(*2*). Presynaptic KARs regulate neurotransmitter release both at the excitatory and inhibitory synapses, while postsynaptic KARs mediate rapid excitatory neurotransmission along with other iGluRs such as the α-amino-3-hydroxy-5-methyl-4-isoxazolepropionic acid and N-Methyl-d-aspartate receptors (AMPARs and NMDARs)(*1, 3*). Consequently, KARs are involved in multiple brain diseases, including schizophrenia and stroke(*4–6*), making them promising targets for clinical intervention. KARs are tetramers assembled from a pool of five distinct subunits (GluK1–GluK5), and each subunit contains four domains; the amino-terminal domain (ATD), the ligand-binding domain (LBD), the transmembrane domain (TMD), and the carboxy-terminal domain (CTD)(*7*). The LBD is composed of two lobes, D1 and D2, which together form a characteristic clamshell-like conformation. Glutamate binding induces closure of the LBD clamshell, thereby increasing tension on the LBD-TMD linkers. This mechanical force pulls apart the TM3 transmembrane helices and opens the ion channel pore. While low agonist affinity (GluK1–GluK3) subunits can form functional homomeric channels, high agonist affinity (GluK4 and GluK5) subunits require co-assembling with GluK1–3 subunits for proper function and localization(*8, 9*). Thus, their diverse subunit combinations are thought to dictate their biophysical properties and physiological roles, similar to AMPARs(*1, 10*). Pre-and postsynaptic KARs also show distinct subtype composition and function(*4, 8, 11*). Native postsynaptic KARs, particularly those found at mossy fiber synapses, are predominantly heteromeric assemblies composed of low-agonist affinity GluK2 and high-agonist affinity GluK5 subunits(*12–15*). Furthermore, native KAR activity is modulated by multiple factors, including auxiliary proteins(*16–23*), extracellular cations and anions(*24–35*), and post-translational modifications(*36–43*).

The functional contributions of GluK2 and GluK5 are thought to be distinct(*44–46*). Notably, the agonist affinity of GluK5 is over 30-fold higher than that of GluK2(*19, 20, 44, 46, 47*). Consequently, activation of GluK5 by low concentrations of glutamate is sufficient to open the channel of heteromeric GluK2/GluK5 KARs, while glutamate binding to the GluK2 subunit, which requires a much higher concentration of the ligand, induces strong receptor desensitization(*48, 49*). Structurally, the heteromeric receptor adopts a 2:2 stoichiometry that comprises two GluK2 and two GluK5 subunits(*50*), organized as a dimer of heterodimers(*51, 52*). Within this arrangement, the GluK5 subunits occupy the A/C positions, while the GluK2 subunits reside in the B/D positions(*53*). Despite decades of studies, the precise mechanisms by which these distinct subunits cooperate to regulate receptor function, including ligand binding, channel activation and desensitization, and how heteromeric KARs differ from their well-studied homomeric counterparts remain elusive. Crucially, while GluK2/GluK5 structures have been reported(*53*), the detailed structural basis of the heteromeric receptors and the specific interactions between GluK2 and GluK5 required to understand these functional differences are still lacking.

In this study, we determined the structure of the heterotetrameric rat GluK2/GluK5 receptor in the apo, antagonized, and desensitized states using single-particle cryo-electron microscopy (cryo-EM). Our structures revealed that, distinct from their homomeric counterparts, the GluK2/GluK5 receptors adopted a highly compact architecture stabilized by extensive intersubunit interactions. These robust interactions, unique to the heteromer, stabilized the classical dimer-of-dimer conformations of the apo and antagonized states as well as the four-fold conformation of the desensitized state in the presence of glutamate. These results highlight the structural advantages that likely favor the formation of heteromeric complexes in native synapses.

Competitive antagonists are compounds that bind to the orthosteric glutamate site, thereby occluding agonist binding and reducing ion flux. While KAR antagonists have been extensively studied in the past decades(*54*) and numerous analogs developed(*14, 54–59*), very few ligands, whether orthosteric or allosteric, have exhibited high subtype selectivity to date. Dr. David Jane and colleagues previously developed a competitive antagonist, (S)-1-(2-amino-2-carboxyethyl)-3-(2-carboxythiophAhene-3-yl-methyl)-5-methylpyrimidine-2,4-dione (UBP310), based on the AMPAR agonist willardiine(*60*). UBP310 is a GluK1/GluK3-selective antagonist that is also effective at GluK2/GluK5 heteromers, albeit with lower affinity(*46*). Crucially, UBP310 exhibits an exceptional approximately 12,700-fold subtype selectivity for GluK1 (the dissociation constant (Kd): 0.13 μM) over GluK2 (Kd: 1.6 mM)(*60*). We utilized this biochemically and electrophysiologically well-characterized compound(*14, 37, 59–65*) in the present study to gain deeper insight into subtype-specific antagonism and investigated the mechanism of subunit-selective antagonism in GluK2/GluK5 receptors. Through the characterization of affinity-altering mutants, we showed that UBP310 adopts distinct binding modes on low-affinity (GluK1–3) versus high-affinity (GluK4 and GluK5) subunits. These differences dictate the receptor’s global architecture. Specifically, UBP310 occupancy at GluK2 resulted in an incomplete pore closure, whereas targeting GluK5 drove the complex into a fully occluded state. Overall, these findings provide a structural and mechanistic framework for understanding the complex gating cycle and pharmacological modulation of KARs.

## Results

### System to specifically analyze the heteromeric GluK2/GluK5 KAR ion channel functions

Previous studies of recombinant heteromeric KARs typically co-transfected GluK2 and GluK5 subunits using varying DNA ratios (e.g., 1:1 to 1:5). This approach may result in a mixed population of homomeric GluK2 and heteromeric GluK2/GluK5 receptors, thereby complicating the specific assessment of the latter. To overcome this limitation and specifically assess the activity of heteromeric subtypes, we employed a receptor trafficking strategy adapted from a system originally used for controlling the assembly of triheteromeric NMDARs(*66–69*). This system, derived from the G protein-coupled GABA_B_ receptor, relies on two peptide tags, C1 and C2, that contain an RXR endoplasmic reticulum (ER) retention signal. These tags were fused to the GluK2 and GluK5 subunits, respectively (Fig. 1A). When expressed individually, the RXR motifs promote endoplasmic reticulum (ER) retention, thus preventing surface trafficking. However, upon co-assembly of the GluK2 and GluK5 subunits into a heteromeric complex, the C1 and C2 tags, which contain rigid leucine zipper motifs derived from GABA_B1_ and GABA_B2_ subunits (LZ1 and LZ2, respectively), interact through a coiled-coil leucine zipper interface. This interaction effectively masks the RXR retention signals, thereby allowing only the correctly assembled GluK2/GluK5 receptors containing two GluK2 and two GluK5 subunits to be transported to the cell surface for functional analysis (Fig. 1A). This method ensured that our electrophysiological and pharmacological data were derived exclusively from the heteromeric GluK2/GluK5 receptors.

**Figure 1.**
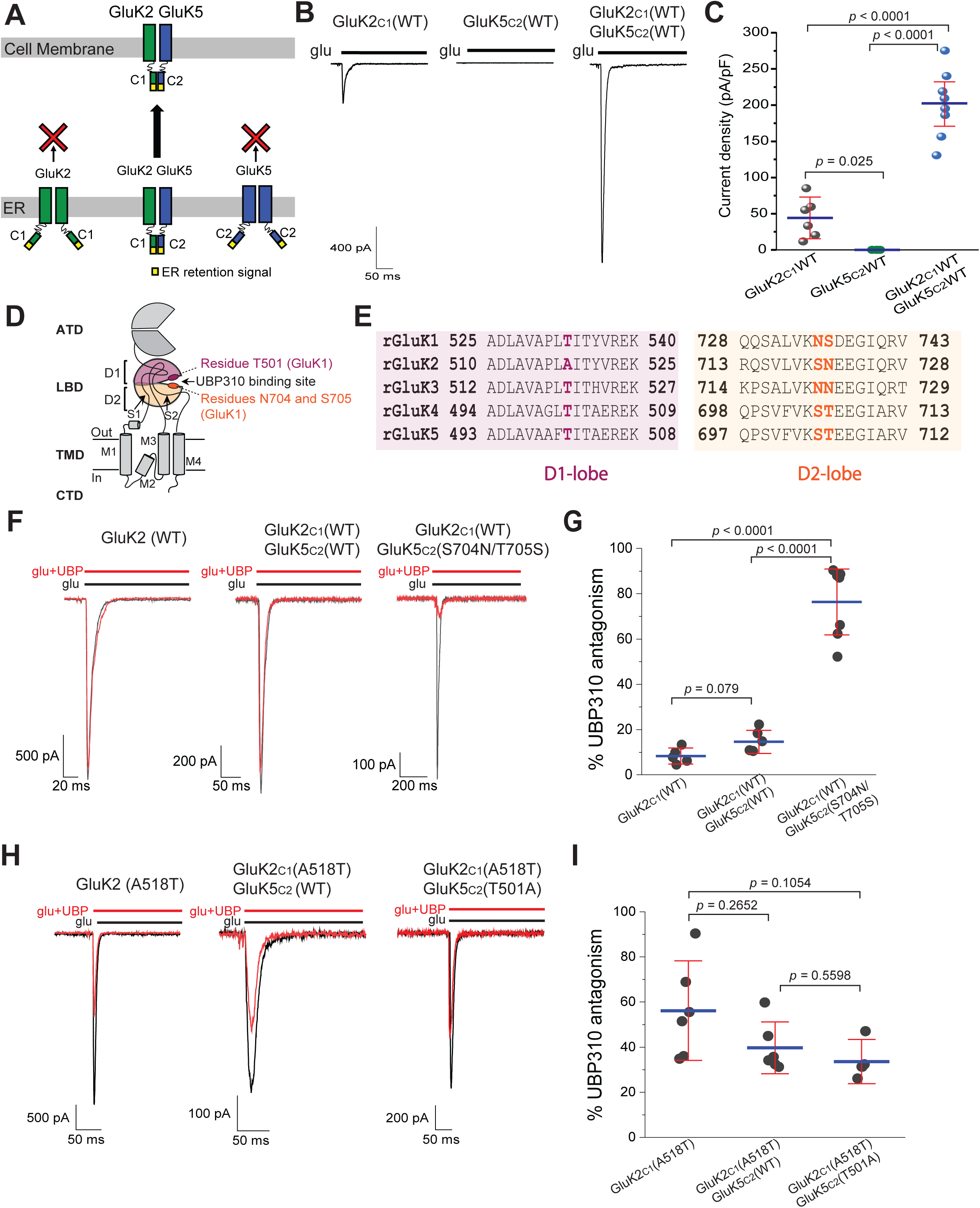
Functional characterization of heteromeric GluK2/GluK5 KARs. **(A)** Strategy for selective surface localization of heteromeric KARs through engineered coiled-coil interactions. The C1 and C2 coiled-coil motifs from the heterodimeric GABA_B1_ and GABA_B2_ receptors, were fused to the C-termini of GluK2 and GluK5, respectively. **(B)** Representative whole-cell voltage-clamp recordings from HEK293T cells expressing GluK2_C1_, left, GluK5_C2_, middle, or heteromeric GluK2_C1_/GluK5_C2_ (right), activated by 10 mM glutamate at-70 mV. Traces are shown at the same scale. **(C)** Current density comparison for the receptor combinations shown in B. **(D)** Schematic of a KAR subunit showing domain organization and residues relevant for UBP310 sensitivity. The ligand-binding domain (LBD) is divided into D1(pink) and D2 (orange) lobes; residue T501 is indicated by a dark pink circle in D1, and residues N704 and S705 (in GluK1) are shown as dark orange circles in D2. **(E)** Sequence alignment of rat KAR subunits (GluK1–GluK5), highlighting regions surrounding the UBP310 binding site, and the positions mutated in GluK2 and GluK5. **(F)** Representative whole-cell currents from GluK2 (left), GluK2_C1_/GluK5_C2_ (middle), and GluK2_C1_/GluK5_C2_S704N/T705S (right) receptors evoked by 10 mM glutamate alone (black) or co-applied with 0.5 mM UBP310 (red). **(G)** Quantification of UBP310 antagonism (% current block) for the receptors shown in F. **(H)** Representative whole-cell current from GluK2 (A518T) (left), GluK2_C1_(A518T)/GluK5_C2_, GluK2_C1_(A518T)/GluK5_C2_ (middle) and GluK2_C1_(A518T)/GluK5_C2_(T501A) (right) receptors evoked by 10 mM glutamate alone (black) or co-applied with 0.5 mM UBP310 (red). **(I)** Quantification of UBP310 antagonism (% current block) for the receptors shown in H. Data are presented as mean ± 95% IC, with individual cells shown as dots. Unless otherwise indicated, statistical analysis was performed using a Welch’s ANOVA followed by Games–Howell post-hoc comparisons. Exact statistics are provided in the Source Data.

To evaluate the efficacy of the C1 and C2 tags, we measured the current amplitude of the rat wild-type GluK2-C1 [GluK2_C1_(WT)] and rat wild-type GluK5-C2 [GluK5_C2_(WT)] when each was expressed individually. GluK5_C2_(WT) showed negligible current, and GluK2_C1_(WT) exhibited only a small current, thus confirming that the transport of C1-or C2-tagged homomers to the plasma membrane is largely blocked and that they are retained in the ER as designed (Fig. 1B). In contrast, co-expression of GluK2_C1_(WT) and GluK5_C2_(WT) yielded current amplitudes exceeding up to ten-fold the homomeric GluK2_C1_ background (Fig. 1B). Comparison of the current density also showed significant differences between the homo-and heteromers [GluK2_C1_(WT): 44.2 ± 27.5 pA/pF, n=6; GluK5_C2_(WT): 0.005 ± 0.009 pA/pF, n=5; and GluK2_C1_(WT)/GluK5_C2_(WT): 201.5 ± 46.0 pA/pF, n=11] (Fig. 1C). These data confirmed the effectiveness of the trafficking strategy for the highly selective functional analysis of heteromeric GluK2/GluK5 receptors.

We next investigated the subtype-specific competitive antagonism of KARs by UBP310, a compound that selectively antagonizes GluK1 and GluK3 but not GluK2(*58, 61, 70*). While UBP310 can also bind to GluK5 within heteromeric KARs(*14, 44, 46*), it has been reported to be significantly less potent at the heteromeric GluK2/GluK5 receptors (IC_50_: 1.3 μM)(*14*) than at homomeric GluK1 (IC_50_: 0.01 μM)(*61, 70–72*). Additionally, previous work by the Molnar and Jane groups demonstrated that the high-affinity binding of UBP310 to GluK1 is primarily mediated by three conserved residues within the LBD: Thr503 on the upper lobe (D1) of the LBD and Asn705 and Ser706 on the lower lobe (D2) of the LBD (Fig. 1D)(*61*). Given that rat GluK5 conserves Thr501 but lacks the other two critical residues on the D2 lobe (Fig. 1E), we introduced the S704N and T705S mutations (corresponding to GluK1 N705/S706) to generate a GluK5 ligand binding pocket mimicking that of GluK1.

To confirm the subtype specificity of UBP310, we first assessed its competitive antagonism on wild-type rat GluK2 (GluK2 WT). Compared to currents activated by 10 mM glutamate alone, the addition of 0.5 mM UBP310 showed only minimal antagonism (8.4 ± 3.4 %, n=5) (Fig. 1F, G). We subsequently co-expressed GluK2 WT with either GluK5 WT or the mutant carrying the S704N/T705S double mutations, tagged with C1 and C2, respectively [GluK2_C1_(WT)/GluK5_C2_(WT) and GluK2_C1_(WT)/GluK5_C2_(S704N/T705S)], and assessed the antagonistic efficacy of UBP310. In contrast to the minimal current reduction observed in homomeric GluK2 WT (8.4 ± 3.4 %, n=5) and heteromeric GluK2_C1_(WT)/GluK5_C2_(WT) to UBP310 (14.6 ± 4.9 % reduced responses, n=6), the heteromeric GluK2_C1_(WT)/GluK5_C2_(S704N/T705S) receptors exhibited a significantly greater reduction in peak current (76.4 ± 16.0 %, n=7) in the presence of 0.5 mM UBP310 (Fig. 1F, G). These results demonstrated that 0.5 mM UBP310 substantially antagonized the GluK2(WT)/GluK5(S704N/T705S) KARs but showed minimal effect on the GluK2 subunit.

To specifically target the GluK2 subunit, we introduced the A518T substitution, an affinity-enhancing mutation previously shown to substantially increase radiolabeled [^3^H]UBP310 binding(*61*). Consistent with the previous finding, UBP310 displayed significantly enhanced competitive antagonism toward the GluK2 A518T mutant by reducing the response by 56.2 ± 21.0 % (n=6), whereas the GluK2 WT receptor remained largely unaffected (Fig. 1F, H, I). Similarly, 0.5 mM UBP310 significantly reduced the responses of the heteromeric GluK2_C1_(A518T)/GluK5_C2_(WT) (40 ± 11 %, n=6) and GluK2_C1_(A518T)/GluK5_C2_(T501A) (33.6 ± 7.9 %, n=5); the latter incorporates the GluK5 T501A mutation, which was engineered to reduce UBP310 binding affinity. These results confirmed the potent competitive antagonism of UBP310 toward these GluK2/GluK5 mutants (Fig. 1H, I). Based on these functional insights, we performed structural analysis using cryo-EM to resolve the corresponding conformational states associated with GluK2/GluK5 activation, desensitization and antagonism.

### Apo GluK2*/GluK5**\*** receptor adopted a dimer-of-dimers architecture stabilized by extensive intersubunit interactions

For the structural studies of the heteromeric KARs, we expressed rat GluK2/GluK5 receptors in HEK293S GnTI⁻ mammalian cells. Due to the low expression and tetramer instability of wild-type GluK2/GluK5 complexes (both of which are full-length and C-terminally truncated), we utilized optimized constructs, which were previously reported by Khanra et al(*53*) that are hereafter referred to as GluK2* and GluK5*. The GluK2* construct underwent RNA editing at position 567 (I to V) and incorporated the C576V and C595S mutations. GluK5* incorporated four cysteine knockout mutations (C559V, C578S, C619I, and C813A) and a GluA2 AMPAR C-terminal tail in place of the original C-terminal domain (ΔS836–E979). Notably, the GluK2*/GluK5* was previously confirmed to form functional heterotetramers(*53*).

To isolate heteromeric complexes from their respective homomeric counterparts, we fused GluK2* and GluK5* to C-terminal twin-strep and 1D4 tags, respectively. We followed the purification protocol we established for determining the apo homomeric GluK2 structure(*29, 38*). Since the previously determined apo GluK2*/GluK5* structure was relatively low resolution (7.5 Å), we first performed single-particle cryo-EM analysis on its apo state. We obtained a substantially lower yield of heteromeric KARs after the three-step purification compared to the high-yields that were achieved for the purified homomeric KARs. To compensate for this, we scaled up purification to 16 L of suspension cell culture and prepared a final protein concentration of a maximum of 1 mg/ml (Fig. S1 A, B). Instead of using 3–4 mg/ml of concentrated protein sample and standard UltrAuFoil R1.2/1.3 300 mesh gold grids as were employed in the previous studies, we utilized the lower protein concentration and prepared cryo-EM grids using copper supports coated with a 2 nm continuous carbon film. Although the overall resolution of the full-length apo-state structure remained at 4.0 Å without applying symmetry, a focused refinement of the LBD layers with applied two-fold symmetry yielded a 3.3 Å reconstruction, thereby enabling a detailed analysis of the LBD layer of the apo GluK2*/GluK5* KARs (Table S1, Fig. S2).

Previous studies from multiple groups, including our own study showed that homomeric apo GluK2 exhibited heterogeneous conformations, ranging from two-fold symmetry to asymmetric arrangements with disrupted LBD dimers(*38, 73*). In contrast, the apo GluK2*/GluK5* dataset solved at 3.3–4 Å primarily resolved into a single dominant class exhibiting an approximately two-fold symmetric conformation (Fig. 2A, B), a result that was consistent with the previously reported apo GluK2*/GluK5* structure at 7.5 Å resolution(*53*). This finding suggested a more stable formation of heteromeric GluK2-GluK5 LBD dimers in the absence of ligands.

**Figure 2.**
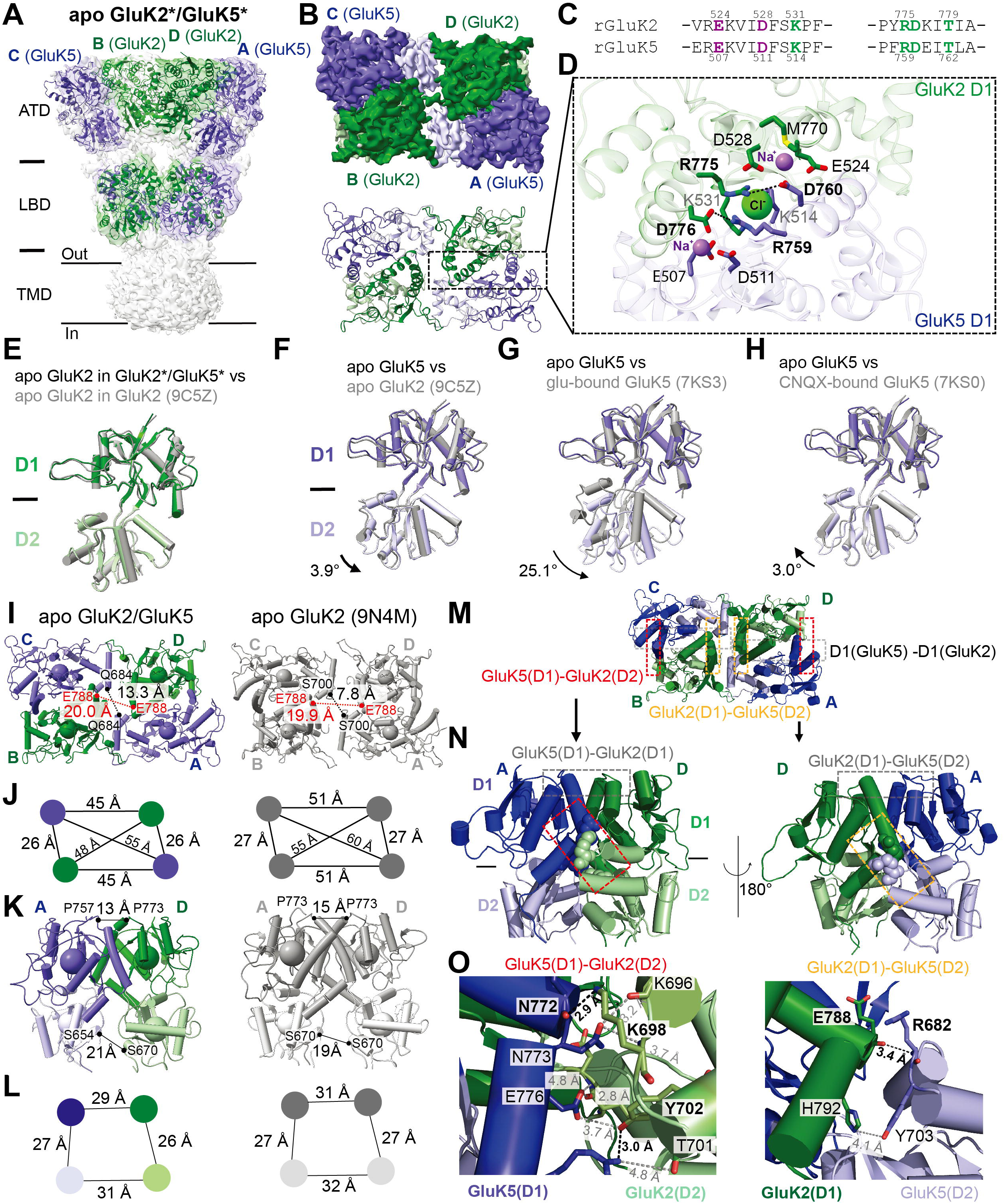
Structural organization of apo GluK2*/GluK5* KARs. **(A)** Cryo-EM density map of apo GluK2*/GluK5* viewed perpendicular to the membrane. Subunits are colored green (GluK2) and purple (GluK5). **(B)** Top view of the LBD layer in the apo GluK2*/GluK5* heteromer. **(C)** Sequence alignment of the Na^+^ and Cl^-^ ion binding sites at the GluK2-GluK5 LBD dimer interface. Conserved residues are highlighted in purple and green, respectively. **(D)** GluK2-GluK5 D1 interface showing key residues mediating intersubunit interactions. **(E)** Superimposition of the GluK2 LBD from the apo heteromeric GluK2*/GluK5* structure (green) with the apo homomeric GluK2 (gray; 9C5Z). **(F–H)** Structural comparisons of GluK5 LBD in the apo GluK2*/GluK5* structure (purple) with glutamate-bound GluK5 (gray; PDB: 7KS3), CNQX-bound GluK5 (PBD: 7KS0), apo GluK2 LBD (PDB: 9C5Z). Rotation angles are indicated. **(I, J)** Intersubunit distances between paired GluK2 LBDs and GluK5 LBDs in the heteromeric GluK2/GluK5 structure (left) versus the homomeric GluK2 structure (right; PDB: 9N4M), viewed from the extracellular side, and distances between the centers of mass (COM) of the D1 and D2 domains. **(K, L)** Arrangement of the GluK2/GluK5 and homomeric GluK2 LBDs. Intersubunit distances are indicated viewed perpendicularly to the membrane, and distances between the COM of the D1 and D2 domains. **(M, N)** Structural overview of D1-D1 and D1-D2 interactions and highlighting the inter-subunit contacts. In addition to the cation-and anion-mediated D1-D1 interface, additional interactions occurred between GluK5 D1 and GluK2 D2 (red) and between GluK2 D1 and GluK5 D2 (yellow). The interactions between N772 (GluK5) and K698 (GluK2) (left) and E788 (GluK2) and R682 (GluK5) (right), which stabilized the assembly, are indicated. The dotted boxes indicate intersubunit interactions between the D1 and D2 lobes: GluK5 D1-GluK2 D2 (red) and GluK2 D1-GluK5 D2 (yellow). **(O)** Close-up view of hydrogen bonds and polar interactions at the GluK2/GluK5 LBD interface.

Similar to the homomeric GluK2 KAR in the apo state(*73, 74*), the apo GluK2*/GluK5* structure showed a direct association between the upper lobes (D1) of the GluK2 and GluK5 LBDs (Fig. 2B–D). Key interactions were formed between Arg775 (GluK2) and Asp760 (GluK5), as well as Asp776 (GluK2) and Arg759 (GluK5) (Fig. 2D). Extensive previous research demonstrated that cation and anion binding stabilizes the D1-D1 association(*24–28*) and thereby regulates the gating kinetics of KARs(*29–35*). Sequence comparison of GluK2 and GluK5 revealed that the residues involved in these cation and anion interactions were conserved (Fig. 2C). In our structure, we observed clear densities corresponding to two sodium ions and one chloride ion, which mediated additional polar interactions between the GluK2 and GluK5 D1 lobes, similar to that seen in the crystal structure of the homomeric GluK2 LBD(*24, 75*) (Fig. 2D, Fig. S3). Consequently, this structure showed conserved D1-D1 interactions, and Na⁺/Cl⁻ binding at the D1-D1 interface in the heteromeric KAR complexes.

The GluK2 LBDs in the apo GluK2*/GluK5* structure were superimposable with the GluK2 LBDs in the homomeric GluK2 structure (PDB: 9C5Z)(*38*) with a root mean square deviation (RMSD) of 0.9 Å (Fig. 2E). Comparison between the apo GluK2 and GluK5 LBDs revealed distinct bi-lobe opening angles: The apo GluK5 LBD was 3.9° more open than the apo GluK2 LBD without ligands (Fig. 2F), thus resulting in an asymmetrical GluK2/GluK5 LBD dimer conformation. As anticipated, the LBD bi-lobe of the apo GluK5 was 25.1° more open than that of the glutamate-bound GluK5 LBD (PDB: 7KS3)(*53*) and 3.0° more closed than the CNQX-bound GluK5 LBD (PDB: 7KS0)(*53*) in the intact GluK2*/GluK5* tetramers (Fig. 2G, H). Beyond the differences in the LBD bi-lobe opening angles, our structural analysis revealed a distinct tetrameric assembly of the apo heteromeric GluK2*/GluK5* and the homomeric GluK2 KARs(*38, 73, 74*). Viewed from the extracellular side, the apo GluK2*/GluK5* complex displayed a more compact conformation (Fig. 2I, J), while the GluK2 D2 and GluK5 D2 lobes were separated in the apo GluK2/GluK5 LBD dimer in the absence of ligands, which was similar to their positioning in the apo homomeric GluK2 (PDB: 9N4M)(*38, 73*) (Fig. 2K, L).

Within the GluK2/GluK5 LBD tetramer, the increased proximity of the GluK2 and GluK5 subunits facilitated a network of inter-subunit hydrogen bonds and polar interactions within the LBD dimer, specifically at the interface between the A(GluK5)-D(GluK2) and B(GluK2)-C(GluK5) pairs (Fig. 2M), in addition to the cation/anion-mediated GluK2 D1-GluK5 D1 interaction (Fig. 2D). When viewed perpendicular to the membrane, the GluK5 D1-GluK2 D2 and GluK2 D1-GluK5 D2 interactions were stabilized by multiple hydrogen bonds and polar contacts on both the exterior and interior surfaces of the GluK2-GluK5 dimers (Fig. 2M–O). These interactions were absent in the homomeric complex.

To assess activation kinetics, we measured the rise time (defined as the time to peak current following glutamate application) for GluK2 WT and GluK2_C1_(WT)/GluK5_C2_(WT) using the receptor trafficking system described above. This comparison showed a non-significant increase of 21 % in *τ*_rise_ (GluK2 *τ*_rise_: 0.9 ± 0.4 ms, n=6, GluK2/GluK5 *τ*_rise_: 1.1 ± 0.3 ms, n=5, *p*= 0.27). Therefore, rather than modulating the rise time, these LBD D1-D2 interactions, which were absent in the apo homomeric GluK2 structure, may uniquely stabilize the tetrameric dimer-of-dimers assembly in heteromeric KARs, thereby complementing the interactions that stabilize individual LBD dimers by reinforcing the dimer-of-dimers conformation of the intact receptor and structurally distinguishing the heteromers from homomeric KARs.

### Structures and distinct binding modes of UBP310 in GluK2*/GluK5* heteromers

To better understand the distinct roles of the high-and low-affinity subunits within the heterotetrameric KARs, we next sought to determine the structure of the heteromeric GluK2*/GluK5* KAR mutants in complex with UBP310. Based on the result of the functional assay described above, we expressed and purified GluK2*/GluK5* KARs that were carrying the GluK5 S704N/T705S double mutation (GluK2*/GluK5*_S704N/T705S_) (Fig. S1C, D), or the combination of GluK2 A518T and GluK5 T501A mutants (GluK2*_A518T_/GluK5*_T501A_) (Fig. S1E, F). 0.5 mM UBP310 was added to the final protein sample, and the mixture was incubated for one hour prior to grid freezing.

The structure of the GluK2*/GluK5*_S704N/T705S_ heteromeric receptors was solved at 4.6 Å (Fig. S4) and adopted an approximately two-fold symmetrical conformation (Fig. 3A, B). Due to the limited resolution of the intact receptor complex, we further performed focused refinement using masks on either the ATD or LBD-TMD layers while applying C2 symmetry. This approach improved the final local resolution of ATD and LBD-TMD to 3.8 Å and 3.9 Å, respectively (Fig. S4, Table S1). Densities corresponding to UBP310 within the GluK5 LBD binding pockets and the TM3 helices were well resolved (Fig. S5A, C).

**Figure 3.**
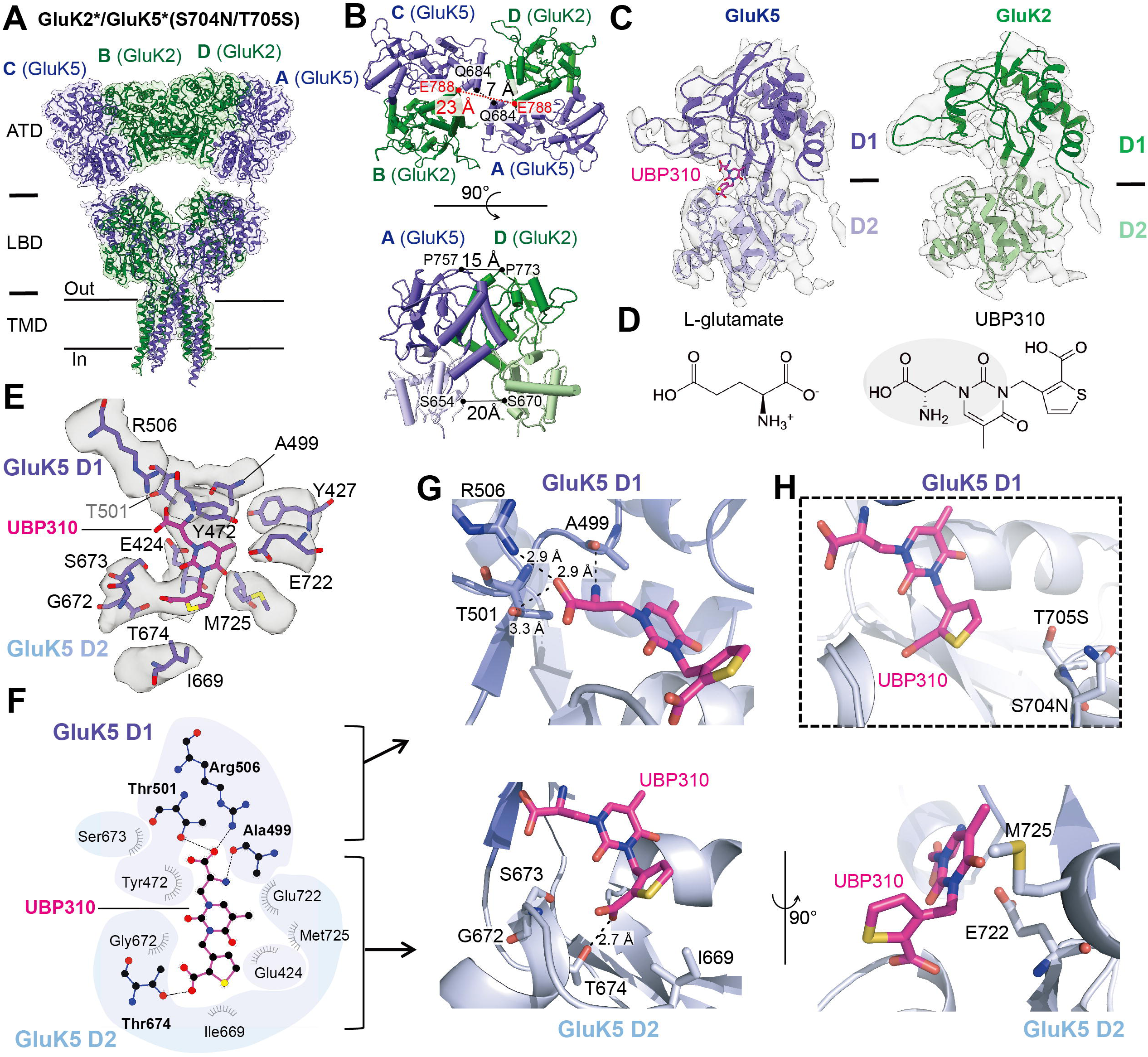
Cryo-EM structure of UBP310-bound GluK2*/GluK5*(S704N/T705S) KARs. **(A)** Cryo-EM density map and model of the UBP310-bound GluK2*/GluK5*(S704N/T705S) heteromer, viewed perpendicular to the membrane. The color scheme is consistent with Figure 1. **(B)** Extracellular view of the LBD layer, highlighting the subunit arrangement within the tetramer. Inter-dimer distances are indicated (top). Side view of a GluK2/GluK5 LBD dimer, showing inter-subunit distances (bottom). **(C)** Comparison of the GluK5 and GluK2 LBDs. The UBP310-bound GluK5 LBD is shown alongside the GluK2 LBD (green) where ligand density was not observed. The panel illustrates the UBP310 binding site at the GluK5 D1-D2 interface. **(D)** Chemical structures of the native agonist L-glutamate and the competitive antagonist UBP310. The gray circle indicates the region of UBP310 that was structurally similar to L-glutamate. **(E)** Cryo-EM density clearly resolving UBP310 bound between the GluK5 LBD D1 and D2 lobes, with key interacting residues labeled. **(F)** Interaction map illustrating the network of hydrogen bonds and polar contacts formed between UBP310 and residues in the GluK5 D1 and D2 lobes. **(G)** Close-up views of the UBP310 binding pocket. Distances between UBP310 and residues forming hydrogen bonds are shown for the GluK5 D1 lobe (top) and the GluK5 D2 lobe (bottom). **(H)** Location of the S704N and T705S mutations introduced in this study, illustrating their position adjacent to the UBP310 binding site without forming direct interactions with the ligand.

Although asymmetrical conformations of antagonist-bound KAR homomers with a disrupted LBD have previously been observed(*37, 76*), even in the presence of auxiliary protein(*77*), we did not identify any classes that exhibited asymmetrical conformation of the heteromeric KARs in the presence of UBP310. In this UBP310-bound heteromeric complex, we also observed polar contacts between the GluK2 and GluK5 LBDs (Fig. S6). Although these inter-subunit interactions were weaker than those in the apo GluK2*/GluK5* structure, they likely contributed to the stability of the tetrameric assembly. These data strongly support that the observed two-fold symmetrical conformation of heteromeric KARs was favored by the robust stability of both the heterodimeric LBD interfaces and the overall heterotetrameric KAR arrangement.

Next, we analyzed the conformational rearrangement within the LBD layer and the detailed UBP310 binding mode on the heteromeric LBDs. We observed that binding of UBP310 brought the two GluK5 LBDs 6.3 Å closer compared to their distance in the apo GluK2*/GluK5* structure, while no substantial changes in the separation between the D1-D1 and D2-D2 lobes were observed (Fig. 3B). As mentioned, clear electron densities corresponding to UBP310 were observed between the D1-D2 lobes of the GluK5 LBDs (Fig. 3C, D, Fig. S5A), which adopted a bi-lobed conformation that was ∼3° more open compared to the GluK5 LBDs in the apo GluK2*/GluK5* structure. In contrast, no discernible ligand densities were detected in the GluK2 LBDs at the same contour level (Fig. 3C, Fig. S7A).

The well-resolved side-chain densities within the ligand binding pocket enabled detailed UBP310-GluK5 interactions in the GluK2*/GluK5*_S704N/T705S_ structure (Fig. 3E). We observed that the D1 lobe of the GluK5 LBD interacted with UBP310 in a manner that closely mimicked its interactions with glutamate (Fig. 3D), similar to the binding mode previously reported in the UBP310-bound GluK1 LBD crystal structures (PDB: 2F34, 2OJT)(*25, 60*). The α-carboxyl group of UBP310 formed hydrogen bonds with the side chain of Arg506 and the main chain oxygen of Thr501, and the amino group formed hydrogen bonds with the side chain of Ala499 (Fig. 3F, G).

In contrast to the conserved interactions between UBP310 and the D1 lobe of GluK5, we observed unique contacts with the D2 lobe that likely contributed to the UBP310 binding affinity. While UBP310 still contacted the D2 lobe, the increased distance primarily resulted in polar interactions rather than the stronger contacts observed in the UBP310-bound GluK1 LBD structure. Specifically, the carboxyl group of UBP310 (attached to the thiophene ring) formed polar interactions with the backbone nitrogen atoms of Ser673, Thr674, and Met725 (Fig. 3E–G). Furthermore, the oxygen atom of the pyrimidinedione ring interacted with the side chain of Glu722. These distances between GluK5 D2 and UBP310 ranged from 3.7 Å to 4.6 Å. Electrophysiological analysis indicated that the double mutations S704N/T705S on GluK5 dramatically increased the binding affinity for UBP310 (Fig. 1F, G). Given that these residues neither formed direct contacts with UBP310 nor interacted with residues directly involved in UBP310 binding (Fig 3H), we propose that they likely function allosterically to stabilize the LBD conformation, thereby enhancing the antagonist affinity.

Next, we determined the structure of the GluK2*_A518T_/GluK5*_T501A_ KARs in the presence of 0.5 mM UBP310 (Fig. 4A). The overall resolution was limited to 7 Å (Fig. S8) primarily due to the low number of particles and preferred orientation issues. However, focused refinement of the LBD and LBD-TMD regions with C2 symmetry application improved the resolutions to 4.8 Å and 5.3 Å, respectively (Fig. S8). Although TM1, TM2, and TM4 were unresolved (Fig. 4A), the relatively clear densities for UBP310 within the GluK2 ligand-binding pockets and all four TM3 helices (Fig. S5B, D) enabled us to assess the ligand binding and the conformation of the ion channel pore.

**Figure 4.**
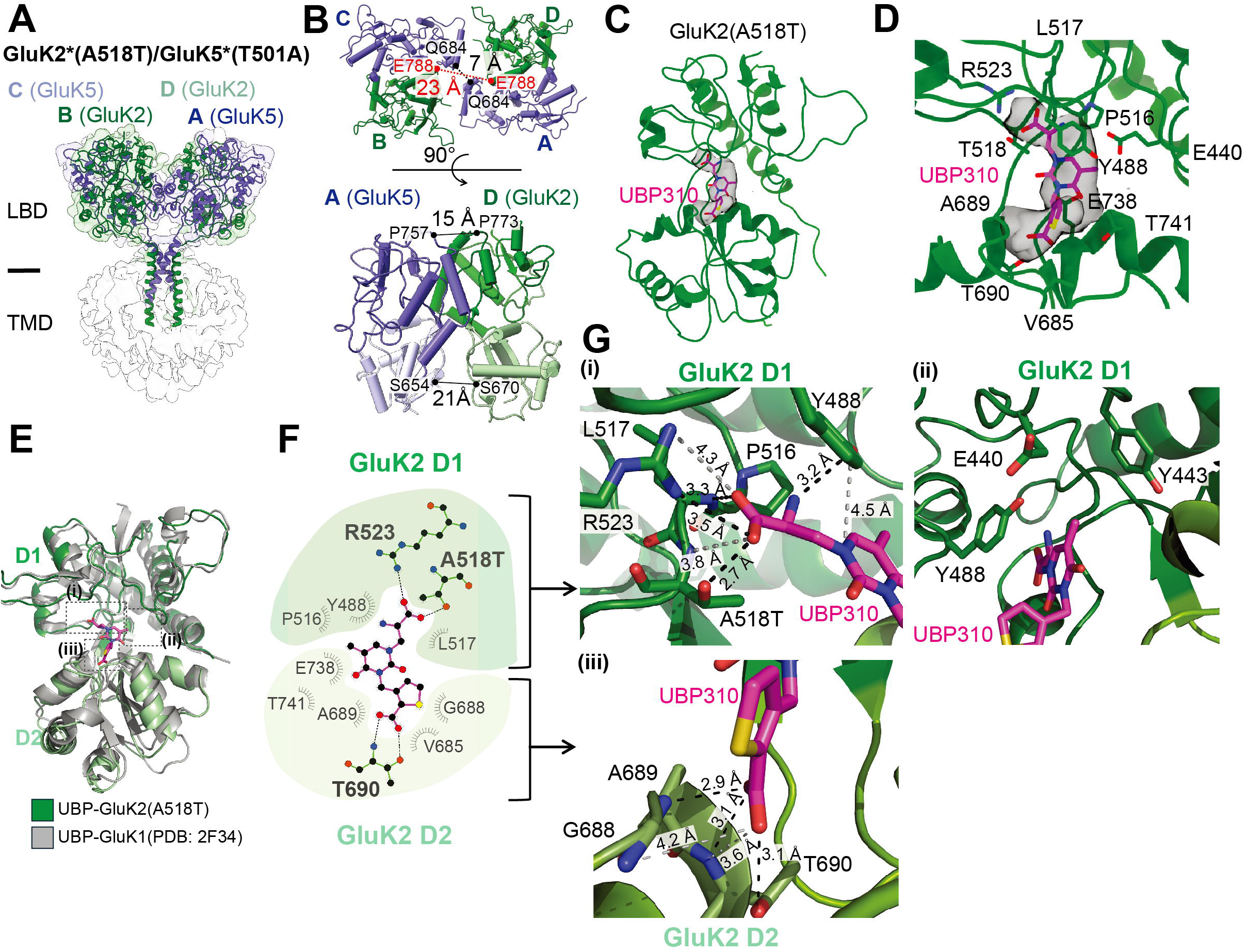
Cryo-EM structure of UBP310-bound GluK2*(A518T)/GluK5*(T501A) KARs. **(A)** Cryo-EM density map and model of the UBP310-bound heteromeric GluK2*(A518T)/GluK5*(T501A) KAR, viewed perpendicular to the membrane. The map resolves the LBD and TM3 helices clearly, whereas the ATD, TM1, TM2, and TM4 regions were poorly resolved due to limited resolution in these domains. **(B)** Extracellular view of the LBD tetramer, highlighting subunit arrangement and inter-dimer distances (top). Side view of a GluK2/GluK5 LBD dimer, showing intersubunit distances (bottom). **(C)** UBP310-bound GluK2 LBD illustrating the ligand position at the D1-D2 interface. The orientation of UBP310 was refined based on the high-resolution GluK1 LBD crystal structure in complex with UBP310 (PDB: 2F34), thereby ensuring accurate positioning of the ligand within the GluK2 binding pocket. **(D)** Close-up view of the UBP310 binding site in the GluK2 ligand binding pocket, showing key interacting residues. **(E)** Superimposition of UBP310-bound GluK2 (A518T) LBD with the UBP310-bound GluK1 structure (PDB: 2F34) highlighting the conformational similarities. **(F)** Interaction map illustrating the network of hydrogen bonds and polar contacts formed between UBP310 and residues in the GluK2 D1 and D2 lobes. **(G)** Close-up views of (i)–(iii) in panel E showing hydrogen bonds between UBP310 and residues in the GluK2 D1and D2 lobes.

The tetrameric LBD assembly in this structure closely resembled that of the GluK2*/GluK5*_S704N/T705S_ complex, including the comparable LBD rolling motion and inter-subunit GluK5 LBD distances (Fig. 4B). While the GluK2*_A518T_/GluK5*_T501A_ structure exhibited clear UBP310 density within the GluK2 LBDs (Fig. 4C, D, Fig. S5B), no corresponding ligand density was observed in the GluK5 LBDs at the same contour level (Fig. S7B). The overall conformation of the UBP310-bound GluK2 LBDs carrying the A518T mutation closely resembled the crystal structure of the UBP310-bound GluK1 LBD (PDB: 2F34)(*60*), with an RMSD of 1.6 Å (Fig. 4E). At a resolution of 4.8 Å, the density was insufficient to unambiguously assign the orientation of the ligand. Therefore, we modeled the UBP310 molecule using the GluK1 LBD structure as a reference. Similar to the GluK1 LBD, both the GluK2 D1 and D2 lobes formed extensive hydrogen bonds and polar interactions with the α-carboxyl group of UBP310 (Fig. 4F, G). The distance between UBP310 and the GluK2 D2 lobe was shorter than the distance between UBP310 and the GluK5 LBD D2 lobe. Consistent with this closer proximity, we observed stronger interactions between the GluK2 D2 lobe and UBP310 compared to the GluK5 D2 lobe and UBP310.

Overall, comparison of the UBP310-bound GluK2* and GluK5* structures revealed distinct binding modes for UBP310 in the high-and low-affinity subunits. This observation directly explains how the unique LBD conformations stabilized by UBP310 influence the compound’s binding affinity and resulting efficacy across these two KAR subunits. To elucidate the subunit-specific antagonism of GluK2 and GluK5, we compared their UBP310-bound structures to define the structural relationships between ligand binding modes, LBD closure, LBD–TMD linker orientation, and ion channel pore architecture.

### Distinct antagonist effects targeting GluK5 or GluK2 subunits

At the LBD layer, UBP310 binding triggered a distinctive approximately 11° rotation of the LBD dimers observed in both UBP310-bound GluK2*/GluK5* mutant structures (Fig. 5A) and the LBD bi-lobe opening (Fig. 5B). This rotation pushed the two GluK2/GluK5 dimers apart by 16.3 Å relative to the apo state, thereby shifting the receptor into more expanded conformations (Fig. 5A). Despite this LBD rotation, the core D1-D1 interface within the LBD dimer remained intact in the presence of UBP310. In the GluK2*/GluK5*_S704N/T705S_ structure, the inter-subunit D1-D1 interface was stabilized by the conserved salt bridges Arg775 (GluK2)-Asp760 (GluK5) and Asp776 (GluK2)-Arg759 (GluK5) (Fig. 5C), as observed in the apo GluK2*/GluK5* structure (Fig. 2C–D). Concurrently, the LBD dimer rotation and GluK5 bi-lobe opening brought the G helices of the two GluK5 subunits into closer proximity. This new arrangement was stabilized by an amide bridge between the Gln684 side chains of opposing GluK5 subunits, as well as polar/electrostatic contacts (Lys797[GluK2]-Asn691 [GluK5] and Asp802[GluK2]-Tyr692 [GluK5]) at the LBD dimer interface (Fig. 5C). While the inter-GluK5 amide bridge was preserved in the GluK2*_A518T_/GluK5*_T501A_ complex, these polar interactions were absent, which likely reflected the reduced degree of opening of the GluK5 LBDs in this mutant.

**Figure 5.**
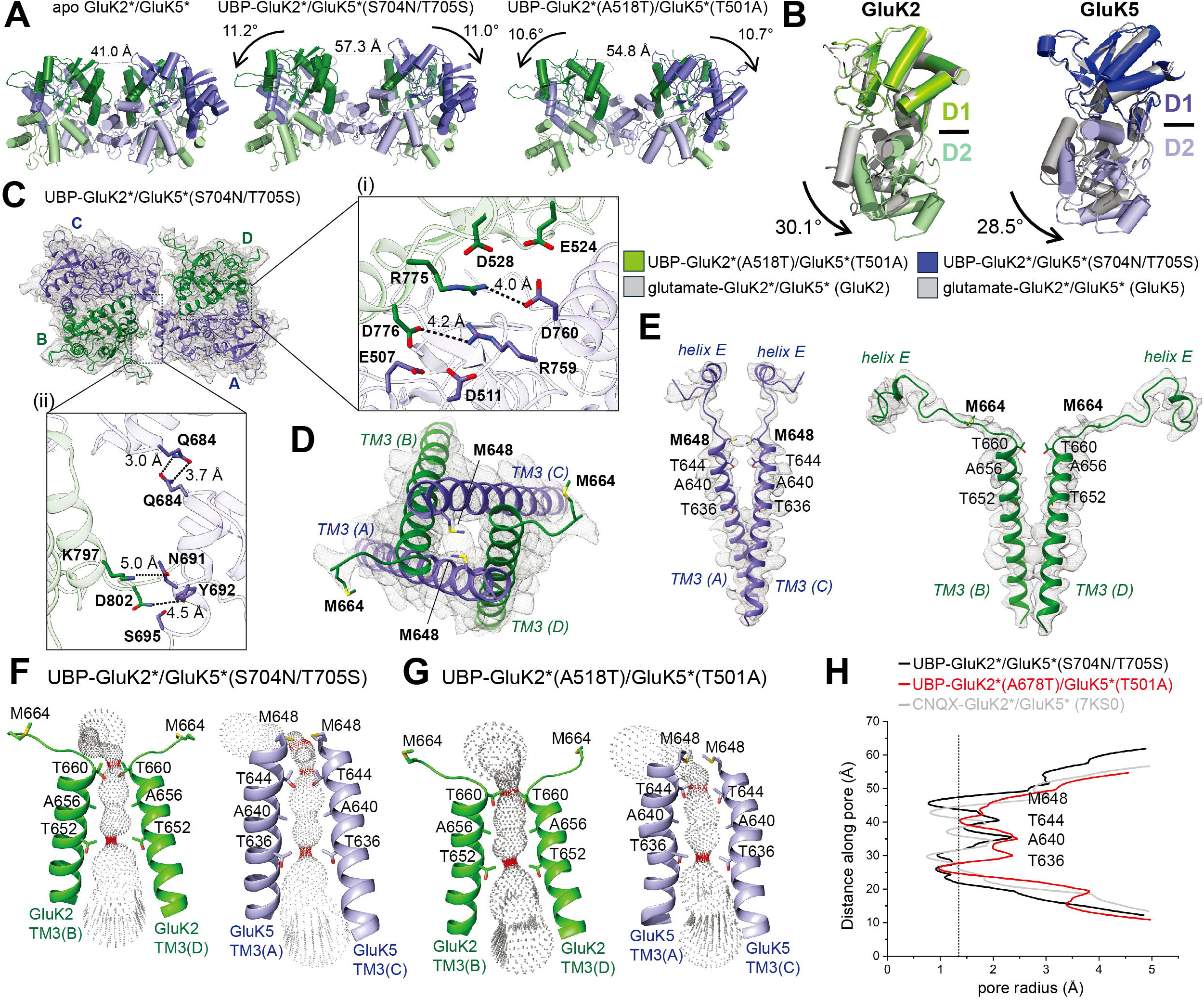
Conformational changes upon UBP310 binding to the GluK2*/GluK5* KARs. **(A)** Comparison of the LBD arrangements and inter-LBD dimer distances in the apo and UBP310-bound GluK2*/GluK5* structures, viewed perpendicular to the membrane. Distances between paired LBDs are indicated. Subunits are colored green (GluK2) and purple (GluK5). **(B)** Superimposition of the UBP310-(colored) and glutamate-bound (gray) GluK2 and GluK5 LBDs highlighting bi-lobe opening. **(C)** Cryo-EM map and atomic model of the LBD layer for the UBP310-bound GluK2*/GluK5*(S704N/T705S) KARs, viewed from the top. Insets highlight the key inter-subunit interactions that stabilized the antagonized state: **(i)** the D1-D1 interface between GluK2 (chain D) and GluK5 (chain A) featuring salt bridges between R775/D760 and D776/R759; **(ii)** the LBD dimer interface between GluK5-GluK5 and GluK2-GluK5 involving hydrogen bonds and polar contacts. **(D)** Cross-sectional view of the transmembrane domain illustrating the closed ion channel pore, occluded by the side chains of Met648 from GluK5 subunits. **(E)** Cryo-EM map and model of helix E and TM3 helix of the GluK2 and GluK5 subunits, showing the positions of key pore-lining residues. **(F, G)** Comparison of pore architecture in the two UBP310-bound GluK2*/GluK5* structures viewed along the channel axis. **(H)** Pore radius profiles calculated for the two UBP310-bound GluK2/GluK5 mutant structures [GluK2*/GluK5*(S704N/T705S) and GluK2*(A518T)/GluK5*(T501A)] and the CNQX-bound GluK2*/GluK5* structure (PDB: 7KS0). The vertical dashed line indicates the radius of a water molecule: 1.4 Å.

Ion channel gating in iGluRs is primarily governed by the ligand-induced opening and closing of the LBDs. A comparison of the UBP310-bound GluK2 and GluK5 LBDs with their apo counterparts revealed no or very small changes in the bi-lobe opening angles (Fig. S7C, D). We therefore compared these LBDs with the fully closed, glutamate-bound GluK2*/GluK5* structures. Comparison of the GluK2 LBD bi-lobe opening angles in the apo GluK2*/GluK5*, UBP310-bound GluK2*/GluK5*_S704N/T705S_, and GluK2*_A518T_/GluK5*_T501A_ structures relative to the glutamate-bound GluK2, yielded values of 23.8°, 25.1°, and 30.1° more opened, respectively. Notably, the UBP310-bound GluK2_A518T_ LBD exhibited the greatest opening at 30.1°, which was 6.3° more open than the apo GluK2 LBD (Fig. 5B). Similarly, we compared the GluK5 LBD bi-lobe opening angles in the apo GluK2*/GluK5*, UBP310-bound GluK2*/GluK5*_S704N/T705S_, and GluK2*_A518T_/GluK5*_T501A_ structures, relative to the fully closed, glutamate-bound GluK5 LBD. The resulting opening angles were 25.2°, 28.5°, and 23.2°, respectively. Because the apo GluK5 LBD (25.2°) adopted a conformation ∼4° more open than the apo GluK2 LBD (23.8°) (Fig. 2F), the difference between the apo and the UBP310-bound GluK5 LBDs was smaller (3.3°) than that observed for GluK2 (6.3°). Nevertheless, the UBP310-bound GluK5_S704N/T705S_ LBD exhibited the greatest opening at 28.5° (Fig. 5B).

These comparisons confirmed that UBP310 binding induces bi-lobe opening in both the GluK2 and GluK5 LBDs, although the magnitude of this structural change differed between subunits, thereby reflecting the distinct UBP310 binding modes. Structural analysis of the UBP310 binding modes revealed distinct ligand orientations and interactions within the GluK2 and GluK5 binding pockets (Fig. S10A–C). While the interacting residues remained largely conserved, the sequence differences between the GluK2 and GluK5 LBDs differences were dictated by the degree of lobe closure and the local electrostatic environment. (Fig. S10D, E).

Despite the absence of clear electron density for the UBP310 in the ligand binding pocket of GluK2 (Fig. S7A), the GluK2 LBD in the GluK2*/GluK5*_S704N/T705S_ structure exhibited an opening angle greater than that of the apo GluK2 LBDs. This observation indicated that the averaged structural data likely included a minority population of GluK2*/GluK5* heteromers where UBP310 was bound to at least one GluK2 subunit, which was consistent with patch-clamp data demonstrating the ∼8% antagonism of homomeric GluK2. This analysis led us to conclude that these structures represented intermediate conformational states, likely resulting from an averaged population of heteromers. In this population, one subunit is strongly bound to UBP310, while the other may be occupied by a weaker or partial binding event, rather than reflecting a state defined by complete subtype selectivity. Nevertheless, this structural analysis allowed us to analyze conformational preferences by comparing these two states.

While the overall tetrameric assembly of the UBP310-bound GluK2*/GluK5*_S704N/T705S_ and GluK2*_A518T_/GluK5*_T501A_ structures was similar (Fig. S7E), we observed small local conformational differences in the GluK5 LBDs, particularly in helices F and I and their connected β-sheets (Fig. S7F). As described above, four residues contributed to UBP310 binding: Ser673 and Thr674 on helix F, and Glu722 and Met725 on helix I (or the adjacent linker within the GluK5 LBD D2 lobe) (Fig. 3G). Helix F was directly linked to helix E, the structural pivot controlling LBD–TM3 linker tension, while helix I could modulate the orientation/position of helix E via a hydrogen bond network between adjacent β-sheets (Fig. S7G). UBP310 binding therefore reoriented the D2 lobes and helix E. In contrast to the compact apo GluK2*/GluK5* receptor, UBP310 induced LBD dimer rotation and D1-D2 cleft opening, which stabilized the heterotetramers in an expanded conformation (Fig. 5A, Fig. S7H). Notably, this arrangement mimicked the more extended structure of the apo homomeric GluK2 receptor (Fig. S7I–K).

This conformation positioned the Met648 of the GluK5 subunits toward the apex of the ion channel, thereby effectively blocking ion entry like a lid (Fig. 5D, E). This state was stabilized by the Gln684 amide bridge and the polar interactions resulting from the GluK5 LBD opening, as described above (Fig. 5C). In the UBP310-bound GluK2*_A518T_/GluK5*_T501A_ complex, the Met648 residues remained but were slightly displaced laterally due to incomplete GluK5 LBD opening in the absence of UBP310 binding, which generated two restriction sites at Thr660(GluK2)/Thr644(GluK5) and Thr652(GluK2)/Thr636(GluK5). In contrast, the UBP310-bound GluK2*/GluK5*_S704N/T705S_ structure showed an additional restriction site at Met648 (GluK5), a conformation similar to the GluK2/GluK5 complex bound to the non-selective antagonist CNQX (PDB: 7KS0)(*53*) (Fig. 5F-H). This analysis indicated that antagonizing GluK5 may more effectively stabilize the closed channel conformation. In summary, our structural analysis supported the functional observation that antagonizing subunits in the A/C positions (e.g., GluK5 in the heteromeric GluK2/GluK5 complex) may be a more effective strategy for stabilizing the non-conducting state.

### Structure of the GluK2/GluK5 in complex with glutamate

Finally, we determined the structure of the glutamate-bound GluK2*/GluK5* KARs in the desensitized state at 3.4 Å without imposing symmetry (C1) (Fig. S9). The complex adopted a four-fold asymmetric conformation that was consistent with the previously reported desensitized GluK2*/GluK5* structure(*53*), with an RMSD of 1.3 Å. Unlike homomeric KARs in the desensitized conformation, which typically show multiple distinct desensitized classes(*38, 78*), the desensitized heteromeric GluK2*/GluK5* also adopted essentially a single, dominant conformation with only minor local differences observed in two minor classes (Fig. S9). This homogeneity suggested that the GluK2*/GluK5* heteromers may have been more conformationally stable than their homomeric counterparts. Consistent with previous reports(*79*), the ATD layer exhibited minimal conformational changes across the apo, antagonized, and desensitized states (Fig. S11). Although the distance between the COM of the ATD and LBD layer in the desensitized state was 7 Å shorter than the apo and antagonized states, this did not result in any new interactions between the ATD and LBD layers. To specifically examine the GluK2-GluK5 interactions within the LBD layer and the ion channel architecture of the TMD layer in the desensitized GluK2*/GluK5* structure, we further performed local refinement of the LBD-TMD layers and achieved a final resolution of 4.0 Å after imposing C2 symmetry (Fig. S9). The composite map, generated from focused ATD and LBD-TMD refinements, showed a highly symmetrical overall conformation (Fig. 6A, B). Our structure allowed us to identify major structural differences between desensitized homo-and heteromeric KARs and to analyze the detailed inter-subunit interactions.

**Figure 6.**
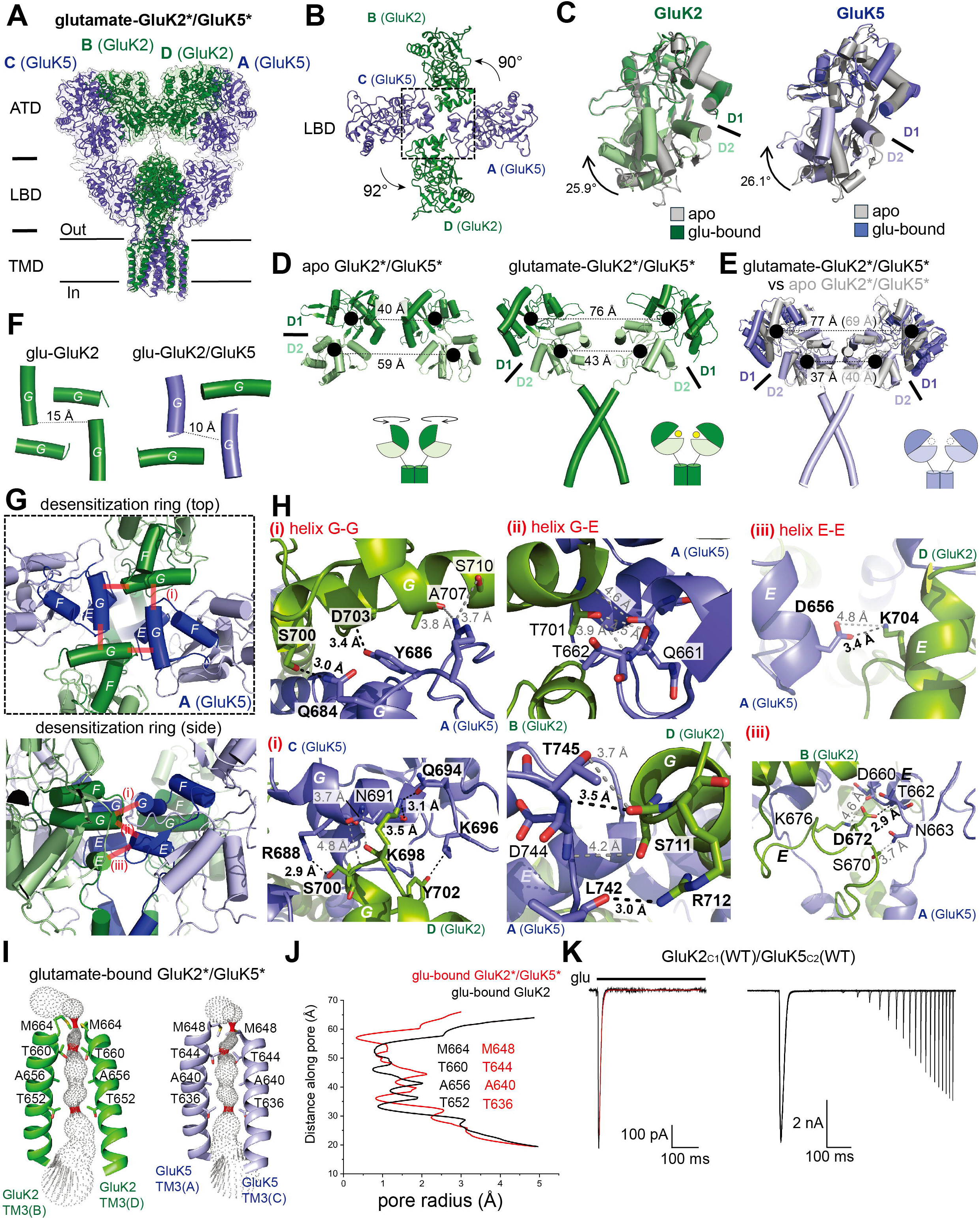
Cryo-EM structure of glutamate-bound GluK2*/GluK5* KARs. **(A)** The glutamate-bound GluK2*/GluK5* KARs in the desensitized state. **(B)** Top-down view of the tetrameric LBD assembly in the glutamate-bound state. Rotation angles associated with the desensitization are indicated. **(C)** Superimposition of the GluK2 and GluK5 LBDs in the apo (gray) and glutamate-bound (colored) GluK2/GluK5. The degree of LBD closure is indicated by arrows. **(D)** Comparison of GluK2 LBD separation in the apo and glutamate-bound states. Distances between the COMs of the D1 and D2 lobes are indicated. **(E)** Superimposition of the apo and glutamate-bound GluK5 LBDs showing minimal overall movement except for D1 and D2 lobe closure upon agonist binding. **(F)** Top-down view of helix G forming the desensitization ring in the glutamate-bound homomeric and heteromeric KARs. **(G)** The desensitization ring in the glutamate-bound GluK2*/GluK5* receptor, viewed parallel (top) or perpendicular (bottom) to the membrane, illustrates the desensitization ring and highlights the inter-subunit hydrogen bonds that stabilize the ring formation. **(H)** Detailed views of the inter-subunit interactions within the desensitization ring stabilizing the desensitized state: (i) the helix G-helix G interface between GluK5 and GluK2, (ii) the helix G-helix E interface, and (iii) the helix E-helix E interface. The key residues involved in the hydrogen bonds and salt bridges are labeled. **(I)** Ion channel pore profiles for the GluK2 and GluK5 subunits in the glutamate-bound state. The pore radius was calculated using HOLE. **(J)** Pore radius profiles calculated for the glutamate-bound GluK2*/GluK5* heteromeric receptor, demonstrating a pore architecture similar to that observed in the glutamate-bound homomeric GluK2 receptors. **(K)** Representative whole-cell currents recorded from HEK293T cells expressing GluK2_C1_/GluK5_C2_ receptors in response to 10 mM glutamate application. These currents were used to calculate desensitization and recovery kinetics.

Although ligand density was not clearly resolved, the LBD bi-lobes of the GluK2 and GluK5 were significantly more closed by 25.9° and 26.1°, respectively, compared to the apo GluK2/GluK5 LBDs (Fig. 6C). This substantial closure strongly suggested full glutamate occupancy on all subunits. When viewed perpendicular to the membrane, the distances between the COM of the D1 domains of the GluK2 and GluK5 LBDs in the desensitized state were nearly identical (∼77 Å), while the distances between the D2 COMs differed slightly, measuring 43 Å (GluK2) and 37 Å (GluK5), respectively (Fig. 6D, E). These matching D1-D1 distances and geometrically similar overall conformations stabilized the tetrameric assembly in the highly symmetric four-fold arrangement. In contrast to the large conformational changes observed in GluK2 (Fig. 6B, D), the comparison of the apo and glutamate-bound GluK2/GluK5 structures revealed that GluK5 underwent minimal structural rearrangement upon desensitization, with changes limited solely to the LBD bi-lobe closure relative to the apo state (Fig. 6E).

To understand the mechanism by which the heterotetramers enhanced structural stability, we further analyzed the LBD layer. It has been shown that KARs characteristically form a ring-like structure composed of helices E and G at the base of the LBDs upon desensitization(*38, 52, 53*). Crucially, when viewed from the extracellular side, the distance between the two GluK5 G helices was maximum 5 Å shorter compared to the corresponding G helices of chains A and C in the homomeric GluK2 structure (Fig. 6F). This reduction in the ring diameter yielded a more compact desensitization ring in the heteromeric GluK2*/GluK5*, thereby resulting in significantly stronger interactions within the ring. Specifically, the interface between the GluK2 and GluK5 subunits featured an increased number of hydrogen bonds and polar interactions compared to those observed in homomeric GluK2(*38, 78*). We observed strong inter-subunit interfaces formed by interactions between helices G-G, helix G-E, and helix E-E (Fig. 6G, H).

Analysis of the ion channel architecture in the desensitized GluK2*/GluK5* structure, specifically the TM3 helices (Fig. S5E), revealed that the pore was tightly closed. It exhibited narrow constrictions at three sites: Met648/Met664, Thr644/Thr660, and Thr636/Thr652 (GluK2/GluK5) (Fig. 6I). Given that the homomeric GluK2 channel already achieved a very tight closure in its desensitized state, the overall pore architecture of the desensitized heteromeric GluK2/GluK5 complex closely resembled that of homomeric GluK2 KAR (Fig. 6J).

To assess how the enhanced structural stability observed in the heteromeric GluK2*/GluK5* KAR influenced receptor kinetics, we assessed two key desensitization-related parameters: the desensitization time constant (*τ*_desensitization_) and the time constant for recovery from desensitization (*τ*_recovery_). We then compared these values with the corresponding kinetics of homomeric GluK2 KAR as reported in our recent study(*38*). We again utilized the heteromeric receptor trafficking system and minimized the background currents originating from the homomeric KARs. We observed that the *τ*_desensitization_ of GluK2_C1_(WT)/GluK5_C2_(WT) was 8.2 ± 2.2 ms (n = 7), which was comparable to that of homomeric GluK2 (7.5 ± 0.3 ms)(*78*) (Fig. 6K). In contrast, the *τ*_recovery_ of GluK2_C1_(WT)/GluK5_C2_(WT) was 1.6 ± 0.7 s (n = 5), significantly slower than that of homomeric GluK2 (716.2 ± 81.0 ms, n=11) (Fig. 6K). Overall, these observations indicated that the desensitized heteromeric KARs were more stable than the homomeric KARs, and this enhanced stability likely underlies the observed prolonged recovery from desensitization.

## Discussion

While native KARs predominantly form heteromers, the detailed molecular mechanism of heteromeric KARs remains elusive due to the lack of high-resolution structures and structures of complexes with subtype-selective ligands. Herein, we reported the structure of the heteromeric GluK2/GluK5 KARs and demonstrated the unique subtype arrangement and inter-subunit contacts that were not conserved in the homomeric KARs.

While the low-affinity subunits GluK1–GluK3 share high sequence similarity (∼96%), the LBDs of the low-affinity subunit GluK2 and high-affinity subunit GluK5 exhibit only ∼78% identity, thereby implying potential differences in conformation. Indeed, our apo GluK2*/GluK5* structure revealed a unique, compact assembly with enhanced conformational stability that distinguished it from homomeric receptors. We observed anions and cations at the GluK2 D1-GluK5 D1 interfaces for the first time and observed that these ions stabilized the GluK2-GluK5 LBD dimers as previously assessed by molecular dynamics simulation(*27*).

Together with unique intersubunit D1-D2 interactions which are absent in homomeric GluK2 KARs(*38, 73*), the apo heteromeric GluK2/GluK5 receptors adopted a more compact and conformationally homogeneous LBD assembly, in contrast to homomeric KARs, which exist as a structural ensemble of both two-fold symmetrical or asymmetrical conformations. Given that KARs can desensitize without prior channel activation(*26*), stabilization of the resting state in a two-fold symmetric conformation may favor a higher proportion of receptors primed for activation rather than partially desensitized receptors as seen in some homomeric KARs(*38, 73, 76*).

The tendency of heteromers to adopt a compact tetrameric arrangement was also evident in the desensitized receptor conformation. The glutamate-bound GluK2*/GluK5* KARs were stabilized in an approximately four-fold symmetric conformation, accompanied by extensive intersubunit contacts in the lower region of the LBD and forming the well-characterized desensitization ring(*52, 53*). These even more robust interactions in heteromers, which involved an increased number of hydrogen bonds and polar contacts compared to homomers, conferred greater structural rigidity and favored a single dominant conformation rather than multiple desensitized states with distinct LBD rotations, as observed previously in homomeric KARs(*38, 53, 78*). While a previous study showed a slightly faster desensitization rate of heteromeric KARs than homomers when analyzed by outside-out patch-clamp electrophysiology(*27*), our whole-cell patch-clamp data showed no clear difference between the desensitization kinetics of GluK2 and GluK2/GluK5. In contrast, we observed that the recovery from desensitization in GluK2/GluK5 was significantly slower than in the homomeric GluK2 KAR. This implied that, once established, the increased stability of the heteromeric desensitization ring may have slowed desensitization recovery. As NETO auxiliary proteins selectively accelerate the recovery of GluK2/GluK5 complexes, while having no such effect on homomeric GluK2(*20*), elucidating the structural basis of this modulation remains a critical objective.

Two structures of UBP310-bound GluK2*/GluK5* mutants revealed distinct binding modes for UBP310 on the GluK2 (GluK1-like) and GluK5 LBDs that directly influence subtype-specificity and competitive antagonism. Since the specific role of GluK5 in receptor function remains poorly understood due to a lack of subtype-specific compounds, the structural details presented here, including the precise degree of LBD bi-lobe opening and the conformation of key residues within the UBP310 binding pocket, provide a critical template for the rational design of the potent GluK5-specific ligands needed to further elucidate the physiological and pathophysiological roles of this subunit.

While antagonizing individual subunits in heteromeric KARs have previously resulted in partial antagonisms(*64*), in contrast to the complete block seen in NMDA receptors(*80*), we observed that antagonizing GluK5 more effectively blocked the channel. Our structures showed that stabilizing the open LBD conformation of GluK5 (at the A/C positions) induced a specific antagonized state where residue Met648 on the LBD-TM3 linker acted as a lid to block the pore entrance, which was specific to GluK5. In contrast, antagonizing GluK2 at the B/D positions is likely less effective because it lacks this pore-blocking system. In summary, our study provides crucial structural insights into the fundamental differences between homomeric and heteromeric KARs and highlights the distinct functional and structural roles of high-and low-affinity subunits within heterotetrameric complexes.

## Methods

### Plasmid construction

For Cryo-EM studies, the construct used for the structural studies was full-length rat GluK2 (Gene bank Nucleotide: NM_019309, Genebank protein: CAA77778.1, Uniprot code: P42260) and C-terminal domain truncated rat GluK5 (Genebank Nucleotide: NM_031508.2, Genebank protein: CAA77667.1, Uniprot code Q63273). Full-length GluK2 gene was RNA edited (I567V), and the construct contains two mutations of C576V and C595S, and fused with a human rhinovirus (HRV) 3C protease recognition site and a C-terminal twinstrep tag(*38*). The GluK5 was mutated (C559V, C578S, C619I, and C813A), truncated at position 827, and added the GluA2 tail (YKSRAEAKRMK), a HRV 3C protease site, and a 1D4 tag which is described previously(*53*). Both genes were synthesized by Genscript and cloned into the pFW/CMV or pFp10/HSP70 vector (gift from Dr. Hiro Furukawa at Cold Spring Harbor Laboratory)(*80, 81*). For the electrophysiological experiments, we subcloned GluK2 in pCAG/IRES/EGFP (Addgene #119739) and GluK5 in pCAG/IRES/mCherry (Addgene # 45766) vectors.

### Cell culture

For structural studies, the bacmid and baculovirus were generated as previously described(*38, 78*). The P1 and P2 viruses were produced in Sf9 cells (Novagen, 71104) in HyClone Insect cell culture media (Cytiva, SH30280.03). HEK293S GnTI^-^ cells (ATCC, CRL-3022) were grown to a density of 3.2 × 10^6^ cells /ml in FreeStyle 293 medium (Gibco, 12338026) at 37°C and 8% CO_2_ supplemented with 2 % fetal bovine serum. HEK293S GnTI^-^ cells were co-infected with the baculovirus harboring GluK2 and GluK5 at the ratio of 1:1 and incubated at 37°C for 12 hrs. Cells were supplemented with 10 mM sodium butyrate (Sigma Aldrich, 303410) and temperature was shifted to 30°C at 12 hrs post-infection. The cell culture was incubated an additional 60 hrs and harvested by low-speed centrifugation at 4,000 rpm for 20 min and stored at-80°C until use. For electrophysiology, Human embryonic kidney 293T (HEK293T) (ATCC, CRL-3216) cells were cultured in Dulbecco’s modified Eagle medium (DMEM, Corning) supplemented with 10% FBS and 1% Penicillin-streptomycin at 37°C in a 95% O₂–5% CO₂ atmosphere. Cells at 70-80% confluence and passages 10-20 were transfected using TransIT-2020 (Mirus) following the manufacturer’s protocol. Rat GluK2 (Grik2) constructs (WT-C1, A478T-C, cloned into pCAG-IRES-EGFP, Addgene plasmid #119739) and rat GluK5 (Grik5) constructs (WT-C2, S704N-T705S-C2 cloned into pCAG-IRES-mCherry; Addgene plasmid #45766) were co-transfected at a [mass/mass] ratio of 1:3 (GluK2:GluK5), with a total of 3 ug DNA per 35 mm dish. C1/C2 Tags correspond to retention endoplasmic reticulum signal, they were inserted at the C-Terminal region.

**C1:** TGEAAAKEAAAKEAAAKEAAAKAMKTGSSTNNNEEEKSRLLEKENRELEKIIAEKEERVSELRHQLQSRQQLKKTN

**C2:** TGEAAAKEAAAKEAAAKEAAAKAVNQASTSRLEGLQSENHRLRMKITELDKDLEEVTMQLQDTPEKKTN

Fluorescence (EGFP/mCherry) was used to identify transfected cells (double-positive fluorescence). After 16-18 h, cells were dissociated with Accutase (Innovative Cell Technologies), re-plated on 35 mm poly-D-lysine-coated dishes (Neuvitro) and recorded 6-12 h post transfection.

### Electrophysiological recordings

All whole-cell patch-clamp recordings were performed using HEKA EPC10 amplifiers (HEKA Elektronik, Lambrecht, Germany) with thin-wall borosilicate glass pipettes (2–5 MΩ) coated with dental wax to reduce electrical noise. Currents were recorded at a holding potential of −70 mV, with a sampling frequency of 10 kHz, and filtered at 2.6 kHz. The external solution contained (in mM): 145 NaCl, 2.5 KCl, 1.8 CaCl₂, 1 MgCl₂, 5 glucose, and 5 HEPES. The internal pipette solution contained (in mM): 105 NaCl, 20 NaF, 5 Na₄BAPTA, 0.5 CaCl₂, 10 Na₂ATP, and 5 HEPES. The pH and osmotic pressure of the external and internal solutions were adjusted to 7.4 and 300–290 mOsm/kg, respectively. L-glutamate (Sigma Aldrich, 49621) was applied using theta glass tubing mounted on a piezoelectric stack (MXPZT-300 series, Siskiyou) driven by a HEKA EPC10 amplifier. Typical 10–90% rise times were 250–300 µs, measured from junction potentials at the open tip of the patch pipette after recordings. Data acquisition was performed using PULSE software (HEKA Elektronik, Lambrecht, Germany).

## Data analysis

Peak current (I_peak_) was quantified as the maximal inward current during agonist application after baseline subtraction. Percent UBP310 antagonism was calculated as:

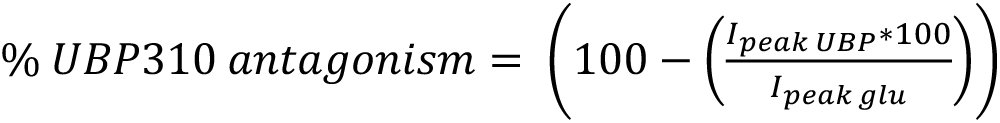

Where *I_peak UBP_* is the response to 500 uM UBP310 (Tocris Bioscience, 3621), and *I_peak glu_* is the control.

**n** denotes independent cells (biological replicates) pooled from at least three independent transfections. Data are presented as mean ± SD unless stated otherwise. Because group variances and sample sizes were unequal, we used Welch’s ANOVA with Games–Howell post-hoc comparisons (two-sided, α = 0.05). Exact P values and effect sizes (ω² for the omnibus test and Hedges’ g for pairwise contrasts) are reported. Analyses were performed in OriginPro 2025 (OriginLab).

Rise time was quantified as the interval required for the current to increase from 10% to 90% of peak amplitude. For a first-order exponential process, this interval corresponds to 2.2*τ*(*4, 82*). Thus, the equivalent activation time constant was estimated as *τ* = (t10–90)/2.2.

### Protein Purification

Cell pellets were resuspended in ice-cold lysis buffer containing 20 mM Tris-Cl pH 8.0, 150 mM NaCl, 0.2 mM PMSF, 20 µg/mL DNase I, and 20 µM DNQX. Cells were lysed by sonication using a Qsonica sonicator at 40% amplitude for a total of 10 minutes (10 s on, 10 s off cycle). The lysate was centrifuged at 7,000 g for 20 minutes to remove cell debris and unbroken cells. The resulting supernatant was subjected to ultracentrifugation at 200,000 g for 30 minutes to pellet cell membranes.

The membrane pellet was homogenized and solubilized by gentle nutation for 2 hours at 4 °C in buffer containing 20 mM Tris-Cl pH 8.0, 150 mM NaCl, 1% (w/v) Dodecyl-β-D-Maltoside (DDM), and 20 µM DNQX. Insoluble material was removed by centrifugation at 200,000 g for 30 minutes. The supernatant was incubated overnight at 4 °C with Twin-Strep-tag affinity resin. The resin-bound protein was extensively washed at room temperature with 100 mL of buffer containing 20 mM Tris-Cl pH 8.0, 300 mM NaCl, 1 mM DNQX, and 0.02% DDM, followed by a slow wash with 200 mL of buffer containing 20 mM Tris-HCl pH 8.0, 300 mM NaCl, and 0.02% DDM at a flow rate of 0.3 ml/min. Protein was eluted in 50 mL of buffer containing 20 mM Tris-Cl pH 8.0, 300 mM NaCl, 0.02% DDM, and 50 mM biotin. The eluted protein was incubated with Rho 1D4 affinity resin for 1 hour at 4 °C. The resin was washed with 50 mL of buffer containing 20 mM Tris-Cl pH 8.0, 500 mM NaCl, and 0.02% DDM at room temperature. Protein was eluted in 50 mL of buffer containing 20 mM Tris-Cl pH 8.0, 300 mM NaCl, 0.02% DDM, and 0.25 mM 1D4 peptide. The eluted protein was incubated overnight with 3C protease to digest and remove EGFP. The sample was then concentrated and injected into a Superose 6 10/300 GL size-exclusion column equilibrated with buffer containing 20 mM Tris-Cl pH 8.0, 150 mM NaCl, and 0.02% DDM. Peak fractions corresponding to tetrameric GluK2/GluK5 were pooled and concentrated to ∼ 1 mg/ml for grid freezing.

### Cryo-EM grids preparation and data acquisition

The cryo-EM grids were prepared in the following manner. Quantifoil Cu 1.2/1.3 400 mesh grids, covered with 2 nm carbon film, were glow-discharged at 20 mA for 20 seconds before sample application.

For the data collection of the apo GluK2*/GluK5*, a droplet of 2.5 µl of protein sample was rapidly applied on grids. For the glutamate-bound GluK2/GluK5 data collection, final concentration of 10 mM L-glutamate was added to the sample and then proteins were pre-incubated for 10 minutes on ice. Volume of 2.5 µL of protein sample was applied on grids. An FEI Vitrobot Mark IV (Thermo Fischer Scientific) was used to plunge-freeze the grids into liquid ethane after sample application at 4 °C and 100% humidity, with a blot time of 2 s and a blot force of 5.

For the data collection of the UBP310-bound and glutamate GluK2*/GluK5*, final concentration of 0.5 mM UBP310 or 10 mM L-glutamate were added to sample and was pre-incubated for 1 hour on ice before applying onto grids.

For the apo GluK2*/GluK5* datasets, all cryo-EM data were collected on a Titan Krios 300kV microscope (Thermo Fisher Scientific) fitted with a GIF-Quantum energy filter (Gatan) with the slit set to 20 e-and a Gatan K3 Summit direct electron detection camera (Gatan). 6,000 and 12,265 movies were collected from 0° and 40° tilted specimens, respectively, using serialEM in the super-resolution mode with a physical image pixel size of 0.64 and a defocus range of-1.0 to-1.5 µm. For the glutamate-bound GluK2*/GluK5* datasets, 7,153 and 18,831 movies were collected on a Titan Krios 300kV microscope from 0° and 40° tilted specimens, respectively. The total dose of around 40 e^-^/A^2^ was used for each movie across 40 frames.

For the UBP310-bound GluK2*/GluK5*(S704N/T705S) datasets, 6,864 and 16,265 movies were collected from 0° and 40° tilted specimens, respectively, using EPU with a physical image pixel size of 0.91 and a defocus range of-1.0 to-1.4 µm.

For the UBP310-bound GluK2*(A518T)/GluK5*(T501A) datasets, 6,365 and 17,653 movies were collected from 0° and 40° tilted specimens, respectively. Both UBP310-bound data were collected on a Glacios 200Kv microscope with slit set to 10 e-and falcon 4 electron detection camera. The total dose of around 40 e^-^/A^2^ was used for each movie across 42 frames.

### Cryo-EM image processing

Apo GluK2*/GluK5*: Raw movie stacks were motion corrected using Patch Motion Correction in CryoSPARC4. Contrast transfer functions (CTFs) were calculated from the motion-corrected images using Patch CTF estimation in CryoSPARC 4. Unsuitable micrographs were removed by manual inspection. Initial particle selection was performed using the Blob Picker from 2000 micrographs, followed by 2D classification to generate templates. Subsequently, 614,826 particles were picked by Template Picker and extracted. After multiple rounds of 2D classification, 125,893 particles were selected to generate improved templates for Topaz picking. 135,217 particles were selected for initial 3D model generation via ab initio reconstruction. Following several rounds of heterogeneous refinement, 56,821 good particles were selected and re-extracted for Non-Uniform (NU) refinement. This procedure yielded a 4.0 Å full-length map. Subsequent LBD-focused local refinement generated a 3.2 Å map.

Glutamate-Bound GluK2*/GluK5*: Data processing followed the same initial procedure as the apo dataset. After 2D classification from initial picking, 822,812 particles were picked by Template Picker and extracted. 178,417 particles were selected for Topaz template generation. 535,217 particles were used to generate the initial 3D model by ab initio reconstruction. 133,425 good particles were selected and re-extracted for NU-refinement after heterogeneous refinement, yielding a 3.41 Å full-length map. Focused local refinement of the ATD and LBD-TMD regions generated 3.13 Å and 4.0 Å maps, respectively.

UBP310-Bound GluK2*/GluK5*(S704N/T705S): Data processing followed the procedure outlined above. 768,451 particles were picked by Template Picker and extracted, with 195,396 particles selected for Topaz template generation. 655,658 particles were used for initial 3D model generation. Following heterogeneous refinement, 153,265 good particles were selected and re-extracted for NU-refinement, which yielded a 4.63 Å full-length map. Particle subtraction was performed for the ATD region, generating a 3.84 Å map by subsequent NU refinement. Subsequent LBD-TMD focused local refinement produced a 3.94 Å map.

UBP310-Bound GluK2*(A518T)/GluK5*(T501A): Data processing followed the same procedure. 565,812 particles were picked by Template Picker and extracted, and 162,864 particles were selected for Topaz template generation.415,968 particles were used for initial 3D model generation. 75,638 good particles were selected and re-extracted for NU-refinement, yielding a 7.04 Å full-length map. Subsequent LBD and LBD-TMD focused local refinement produced 4.85 Å and 5.33 Å maps, respectively.

### Model building and refinement

The model building for the apo GluK2*/GluK5* was commenced by rigid-body fitting of the individual LBD extracted from the cryo-EM structure (PDB: 9C5Z) and the predicted GluK5 model, and the ATD extracted from the cryo-EM structure (PDB: 7KS0) into the cryo-EM map using USCF Chimera X. Subsequent manual adjustments, including fitting the backbone and side chain of the ATD-LBD linkers into the density, deletion of unresolved side chains, and subtle local alterations for clash minimization, were carried out in Coot.

The model building for the glutamate-bound GluK2*/GluK5* was commenced by rigid-body fitting of the individual LBD extracted from the cryo-EM structure (PDB: 7F57) and the predicted GluK5 model, and the ATD extracted from the cryo-EM structure (PDB: 7KS3) into the cryo-EM map using Chimera X. Subsequent manual adjustments, including fitting the backbone and side chain of the ATD-LBD and LBD-TMD linkers into the density, deletion of unresolved side chains, and subtle local alterations for clash minimization, were carried out in Coot.

The model building for the UBP310-bound GluK2*/GluK5*(S704N/T705S) was commenced by rigid-body fitting of the individual LBD extracted from the cryo-EM structure (PDB: 9C5Z) and the predicted GluK5 model, and the ATD extracted from the cryo-EM structure (PDB: 7KS0) into the cryo-EM map using Chimera X. Subsequent manual adjustments, including fitting the backbone and side chain of the ATD-LBD and LBD-TMD linkers into the density, deletion of unresolved side chains, and subtle local alterations for clash minimization, were carried out in Coot. The UBP310 compound model was extracted from the structure (PDB: 2F34) and manually fit into the corresponding densities in Coot.

The model building for the UBP310-bound GluK2*(A518T)/GluK5*(T501A) was commenced by rigid-body fitting of the individual LBD extracted from the cryo-EM structure (PDB: 9C5Z) and the predicted GluK5 model, and the ATD extracted from the cryo-EM structure (PDB: 7KS0) into the cryo-EM map using Chimera X. Subsequent manual adjustments, including fitting the backbone and side chain of the ATD-LBD and LBD-TMD linkers into the density, deletion of unresolved side chains, and subtle local alterations for clash minimization, were carried out in Coot. The UBP310 compound model was extracted from structure (PDB: 2F34) and manually fit into the corresponding densities in Coot.

All initial models were further refined in real space by Phenix against the corresponding cryo-EM map. All models were validated by comprehensive validation in Phenix.

## Acknowledgment

We thank Dr. Zhongwu Zhou at Cleveland Clinic, for technical assistance with stage tilting during EM data collection. We also thank Drs. Kunpeng Li and Kyle Whiddon at Case Western Reserve University (CWRU) for their cryo-EM technical support (NIH S10OD03243701), the High-Performance Computing Resource CWRU Core facility for their computational support, and Dr. Corey Smith and Dr. Shyue-An Chan at CWRU for assistance with patch-clamp electrophysiology. We are grateful to Dr. Matthias Buck at CWRU for comments on this work.

## Author Contributions

Conceptualization: C.Z., G.S.-C., A.H., and N.T. Methodology: C.Z., G.S.-C., A.H., and N.T. Investigation: C.Z., G.S.-C., A.H., and N.T. Visualization: C.Z., G.S.-C., A.H., and N.T. Resources: C.Z., G.S.-C., and N.T. Funding acquisition: G.S.-C., and N.T. Supervision: N.T. Project administration: C.Z., G.S.-C., and N.T. Data curation: C.Z., G.S.-C., and N.T. Validation: C.Z., G.S.-C., A.H., and N.T. Formal analysis: C.Z., G.S.-C., A.H., and N.T. Writing—original draft: C.Z., G.S.-C., A.H., and N.T Writing—review and editing: C.Z., G.S.-C., A.H., and N.T. Software: C.Z., G.S.-C., and N.T.

## Data, Code, and Materials Availability

All data and code needed to evaluate and reproduce the results in the paper are present in the paper and/or the Supplementary Materials. There are no new materials generated in the study. The cryo-EM density maps and coordinates for GluK2/GluK5 have been deposited in the Electron Microscopy Data Bank under accession codes numbers EMD-73886, EMD-73887, EMD-73888, EMD-73889, EMD-73890, EMD-73891, EMD-73892, EMD-73893, EMD-73894, EMD-73895, EMD-73897, EMD-73898, and in the PDB under accession codes 9Z85, 9Z86, 9Z87, 9Z88, 9Z89, 9Z8A, 9Z8B, 9C8Z, 9Z8D, 9Z8E, 9Z8G, 9Z8H. Requests for the plasmids generated for this research should be submitted to Nami Tajima at.

## List of the supplemental materials

Figure S1: Figure S1.pdf

Figure S2: Figure S2.pdf

Figure S3: Figure S3.pdf

Figure S4: Figure S4.pdf

Figure S5: Figure S5.pdf

Figure S6: Figure S6.pdf

Figure S7: Figure S7.pdf

Figure S8: Figure S8.pdf

Figure S9: Figure S9.pdf

Figure S10: Figure S10.pdf

Figure S11: Figure S11.pdf

Table S1: Table S1.pdf

