## Supplemental figures for "Cryo-EM Structures of Apo, Agonist- and Antagonist-Bound Heteromeric Kainate Receptors"

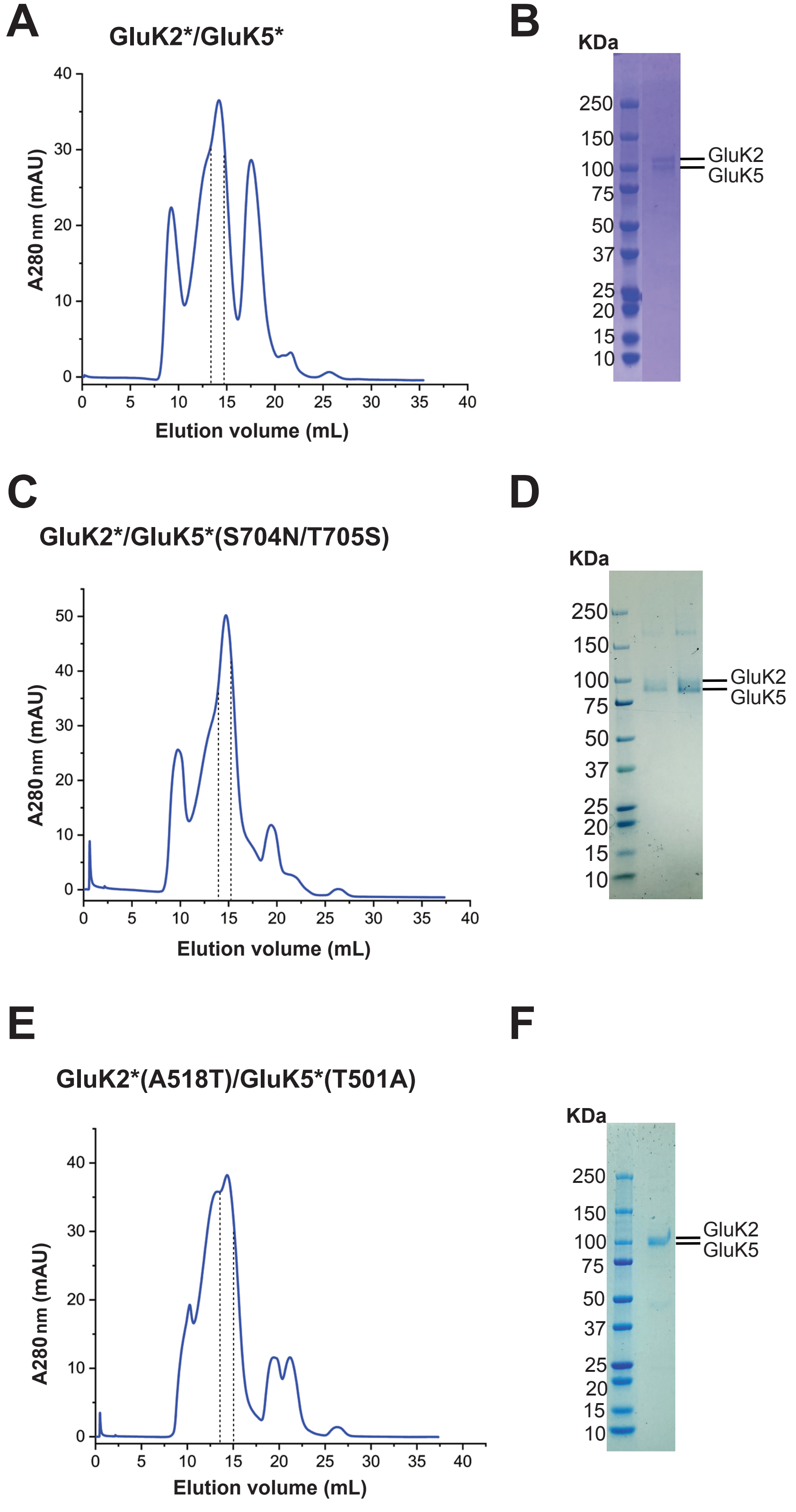


**Figure S1. Size-exclusion chromatography and SDS–PAGE analysis of GluK2*/GluK5* KARs.**
**(A, C, E)** SEC profiles showing peaks corresponding to receptor assembly. Fractions containing the tetrameric GluK2*/GluK5* KARs used for the cryo-EM studies are indicated. **(B, D, F)** SDS-PAGE analysis of the final protein samples used for the cryo-EM studies. The bands corresponding to the GluK2 and GluK5 subunits are labeled.

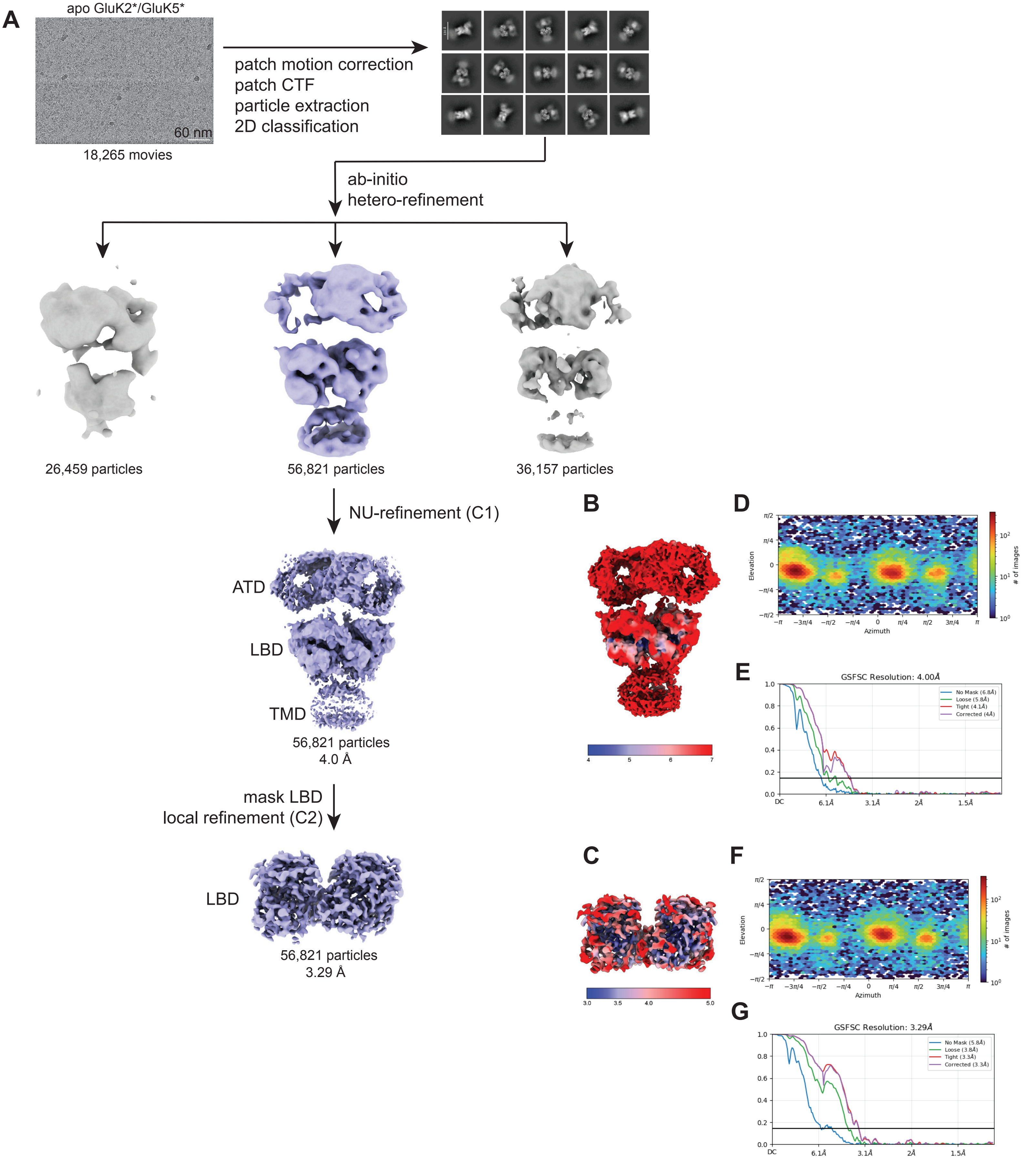


**Figure S2. Cryo-EM data processing workflow for the apo GluK2*/GluK5* KARs.**
**(A)** CryoSPARC-based data processing flowchart. **(B, C)** Local resolution estimation of the cryo-EM maps for full-length and LBD, respectively. **(D, F)** Fourier shell correlation (FSC) curves of half-maps for full-length and LBD. **(E, G)** Particle angular distribution for the final 3D reconstruction for the respective maps.

**A** **apo GluK2*/GluK5* KAR**

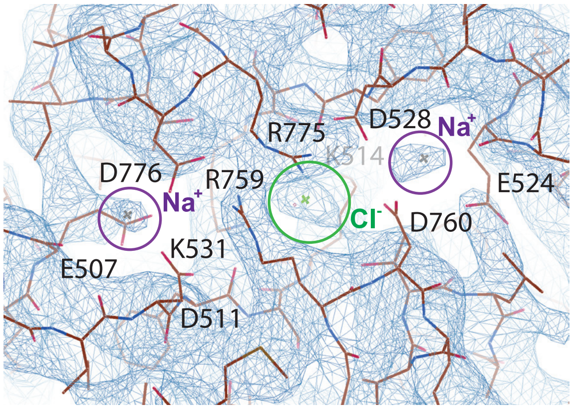


**Figure S3. Ion-mediated interactions at the D1-D1 interface of the GluK2*/GluK5* KARs.**
**(A)** Cryo-EM density map and atomic model showing the D1-D1 interface between the GluK2 and GluK5 subunits in the apo state. Key residues involved in the inter-subunit interactions are labeled, along with the coordinated ions ($Na^+$ and $Cl^-$) that stabilized the interface. The mesh represents the cryo-EM density contoured around the interacting residues.

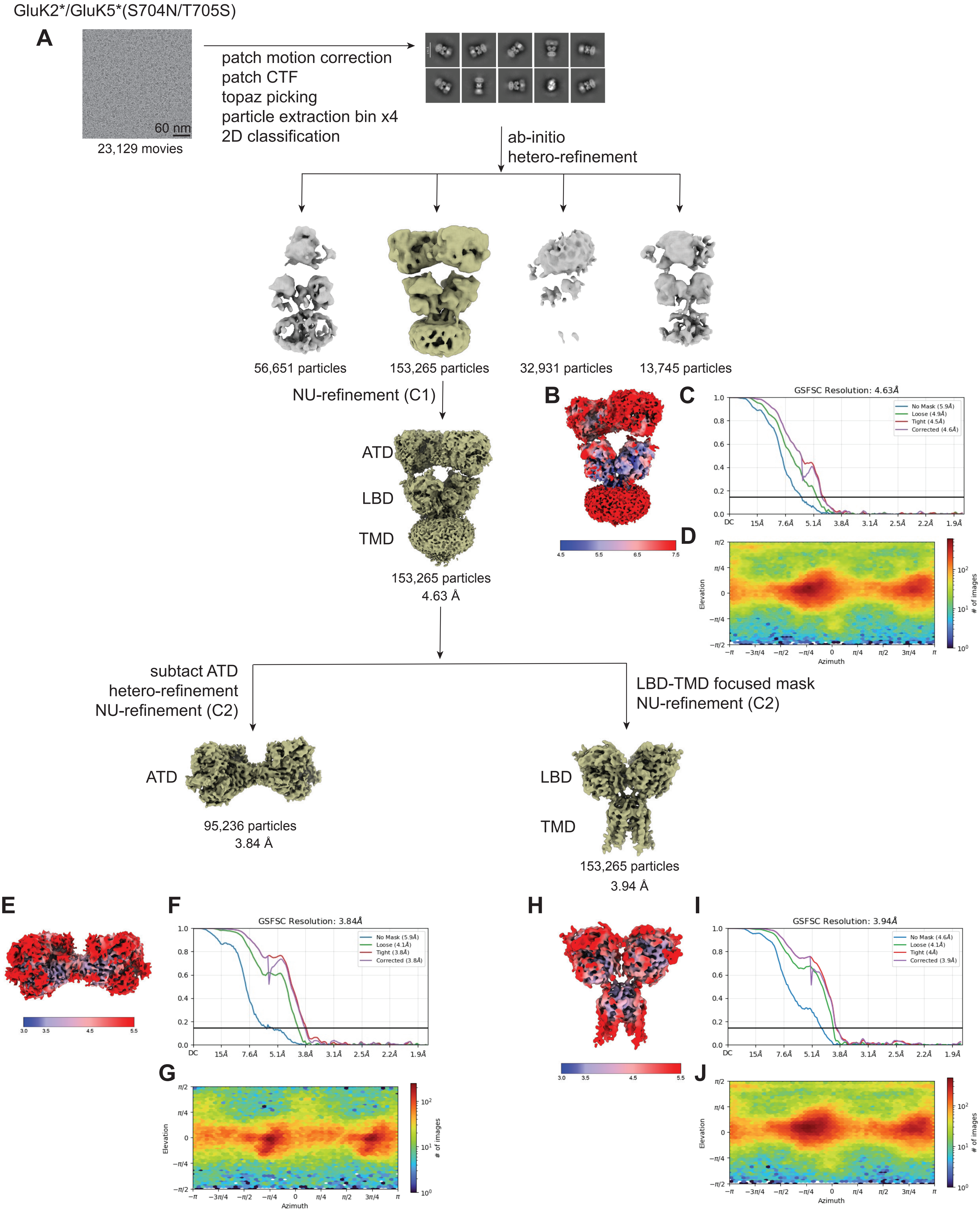
GluK2*/GluK5*(S704N/T705S)
A
60 nm
23,129 movies
patch motion correction
patch CTF
topaz picking
particle extraction bin x4
2D classification
ab-initio
hetero-refinement
56,651 particles
153,265 particles
32,931 particles
13,745 particles
NU-refinement (C1)
ATD
LBD
TMD
153,265 particles
4.63 Å
B
C
GSFSC Resolution: 4.63Å
No Mask (5.9Å)
Loose (4.9Å)
Tight (4.5Å)
Corrected (4.6Å)
D
Elevation
Azimuth
### of images
subtact ATD
hetero-refinement
NU-refinement (C2)
LBD-TMD focused mask
NU-refinement (C2)
ATD
95,236 particles
3.84 Å
LBD
TMD
153,265 particles
3.94 Å
E
F
GSFSC Resolution: 3.84Å
No Mask (5.9Å)
Loose (4.1Å)
Tight (3.8Å)
Corrected (3.8Å)
G
H
I
GSFSC Resolution: 3.94Å
No Mask (4.6Å)
Loose (4.1Å)
Tight (4Å)
Corrected (3.9Å)
J

**Figure S4. Cryo-EM data processing workflow for the UBP310-bound GluK2*/GluK5*(S704N/T705S) KARs.**
**(A)** CryoSPARC-based data processing flowchart. **(B, E, H)** Local resolution estimation of the cryo-EM maps for the respective structures. **(C, F, I)** FSC curves of half- maps for full-length, ATD and LBD, respectively.
**(D, G, J)** Particle angular distribution for the final 3D reconstruction for respective maps.

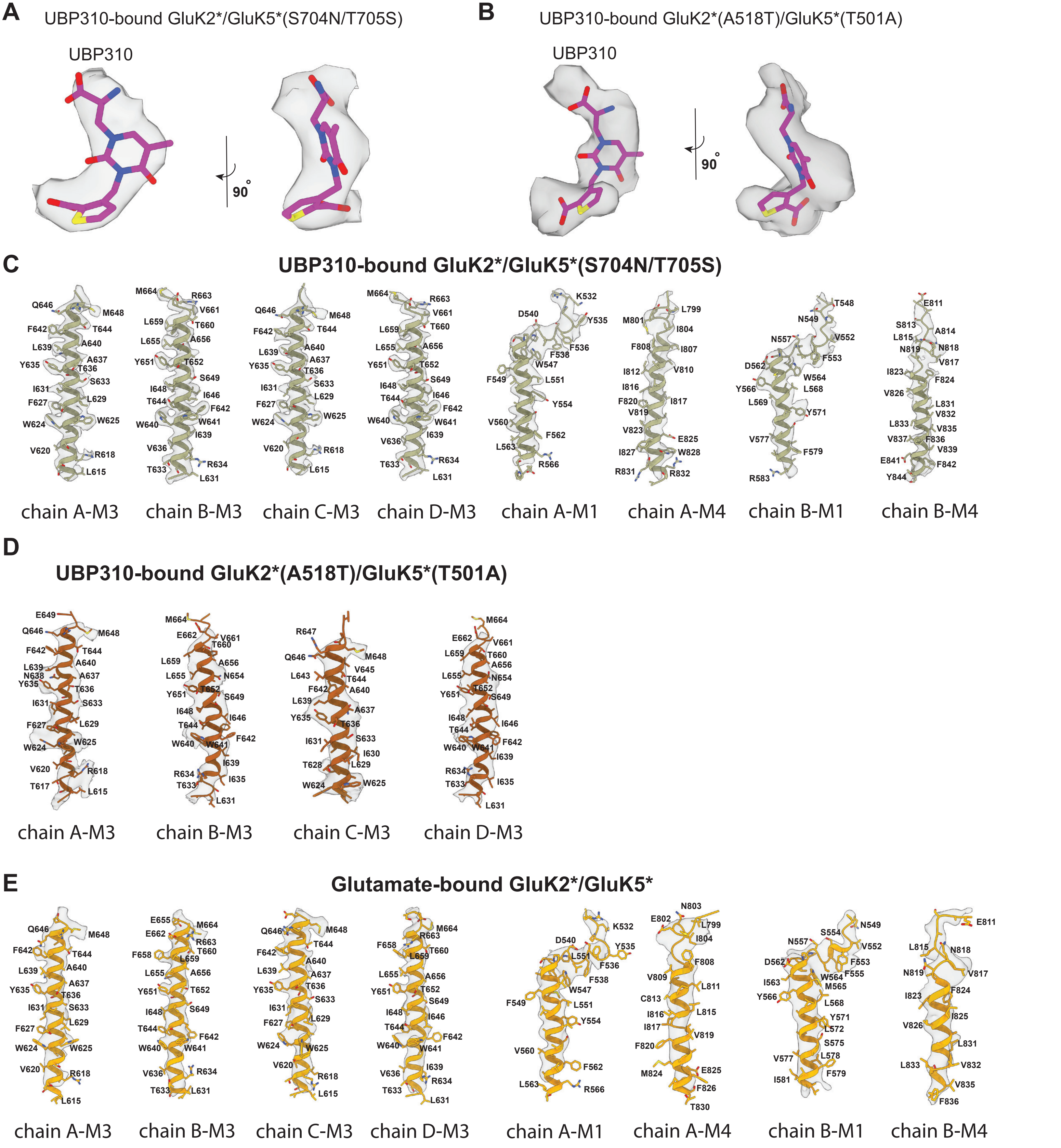


**Figure S5. Cryo-EM densities of the ligands and transmembrane helices.**
**(A)** Cryo-EM density and model of UBP310 bound to the GluK5 subunit in the UBP310-bound GluK2*/GluK5*(S704N/T705S) structure. **(B)** Cryo-EM density and model of UBP310 bound to the GluK2 subunit in the UBP310-bound GluK2*(A518T)/GluK5*(T501A) structure. **(C–E)** Cryo-EM density and model of the transmembrane helices from the UBP310-bound GluK2*/GluK5*(S704N/T705S) (C), UBP310-bound GluK2*(A518T)/GluK5*(T501A) (D), and glutamate-bound GluK2*/GluK5* (E) structures.

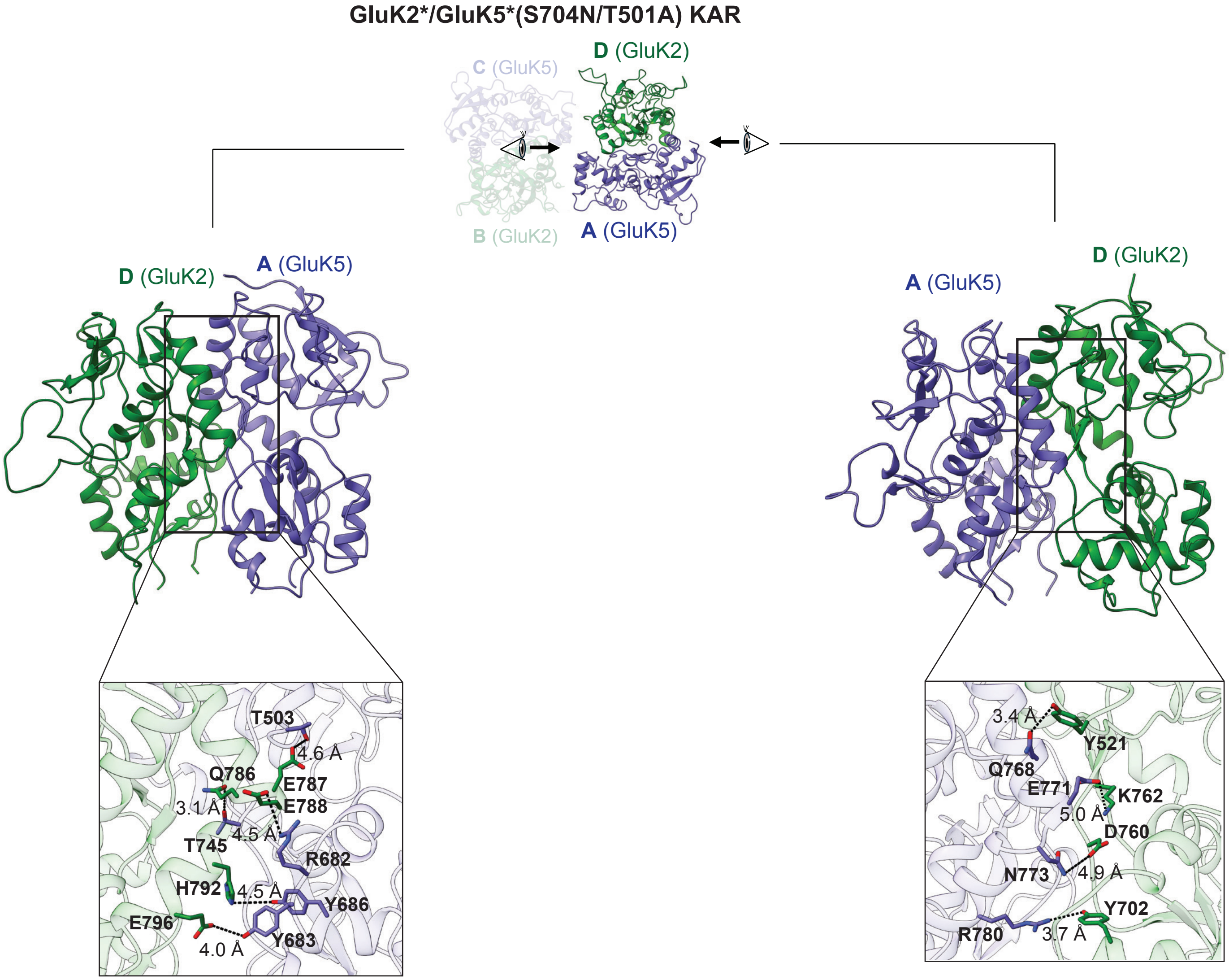


**Figure S6. Inter-dimer LBD contacts in the GluK2*/GluK5*(S704N/T705S) KARs.**
A detailed view of the inter-subunit D1-D1 interface between the GluK2 and GluK5 subunits in the presence of UBP310. Key stabilizing interactions are indicated, including salt bridges and polar contacts that maintained the integrity of the D1-D1 interface despite the overall LBD layer rearrangement.

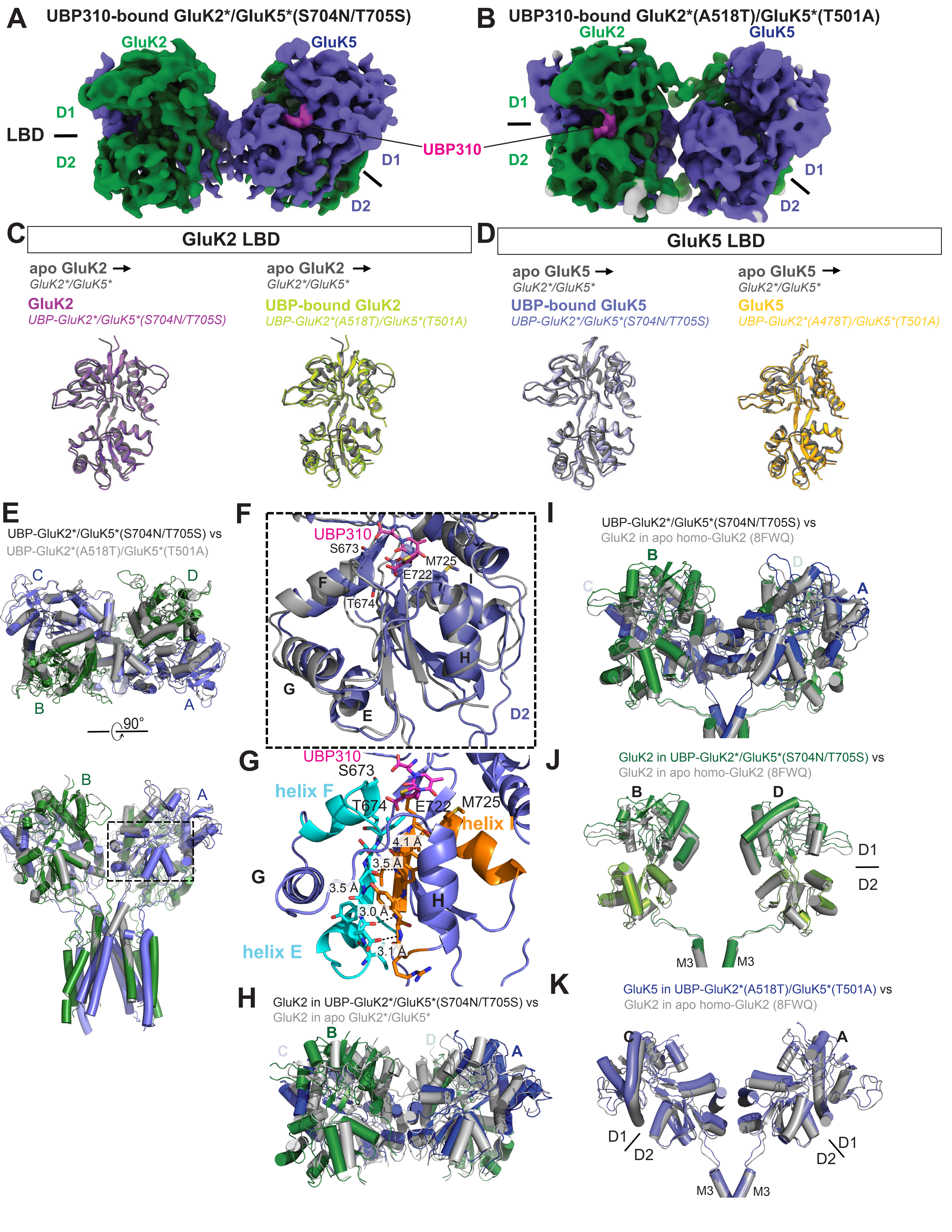
A UBP310-bound GluK2*/GluK5*(S704N/T705S)
B UBP310-bound GluK2*(A518T)/GluK5*(T501A)
GluK2
GluK5
LBD
D1
D2
UBP310
C GluK2 LBD
D GluK5 LBD
apo GluK2 →
GluK2*/GluK5*
GluK2
UBP-GluK2*/GluK5*(S704N/T705S)
UBP-bound GluK2
UBP-GluK2*(A518T)/GluK5*(T501A)
apo GluK5 →
UBP-bound GluK5
GluK5
UBP-GluK2*(A478T)/GluK5*(T501A)
E
UBP-GluK2*/GluK5*(S704N/T705S) vs
UBP-GluK2*(A518T)/GluK5*(T501A)
90°
F
S673
M725
E722
T674
G
helix F
helix E
helix I
4.1 Å
3.5 Å
3.0 Å
3.1 Å
I
UBP-GluK2*/GluK5*(S704N/T705S) vs
GluK2 in apo homo-GluK2 (8FWQ)
J
GluK2 in UBP-GluK2*/GluK5*(S704N/T705S) vs
GluK2 in apo homo-GluK2 (8FWQ)
M3
H
GluK2 in UBP-GluK2*/GluK5*(S704N/T705S) vs
GluK2 in apo GluK2*/GluK5*
K
GluK5 in UBP-GluK2*(A518T)/GluK5*(T501A) vs
GluK2 in apo homo-GluK2 (8FWQ)

**Figure S7. Structural comparison of UBP310-bound GluK2*/GluK5* KAR mutants.**
**(A, B)** Cryo-EM density maps of the UBP310-bound GluK2*/GluK5*(S704N/T705S) and GluK2*(A518T)/GluK5*(T701A) heteromeric receptors. Subunits are color-coded as GluK2 (green) and GluK5 (purple), with resolved UBP310 ligand density indicated in pink. **(C)** Superimposition of GluK2 LBD structures in apo compared with UBP310-bound conformations for both mutant complexes. **(D)** Superimposition of GluK5 LBD structures in apo compared with UBP310-bound conformations for both mutant complexes. **(E)** Superimposition of the full receptor structures of UBP310-bound GluK2*/GluK5*(S704N/T705S) (colored by subunit: green/purple) and UBP310-bound GluK2*(A518T)/GluK5*(T501A) (gray). Top view (top) and side view (bottom) show similar conformation of the two mutant complexes. **(F)** Detailed view of the boxed region in E, highlighting the LBD D2 lobe in the UBP310-bound GluK2*/GluK5*(S704N/T705S) structure. The key residues involved in the interactions with UBP310 are labeled. **(G)** Close-up view of the β-sheets connected to helix F, helix G, and helix I in the UBP310-bound complex. Distances between residues on the β-sheets are indicated to highlight the stabilization of the interface.
**(H, I)** Superimposition of the LBD layer of UBP310-bound GluK2*/GluK5*(S704N/T705S) (green/purple) onto the apo GluK2*/GluK5* structure (gray) (H), or the apo homomeric GluK2 structure (PDB: 5WJQ, gray) (I), illustrating differences in quaternary arrangement. **(J, K)** Comparison of the GluK2 (J) and GluK5 (K) subunit conformations. GluK2 and GluK5 from the UBP310-bound GluK2*/GluK5*(S704N/T705S) complex (green/puple) were superimposed on the GluK2 subunit from the apo homomeric GluK2 (5WJQ, gray).

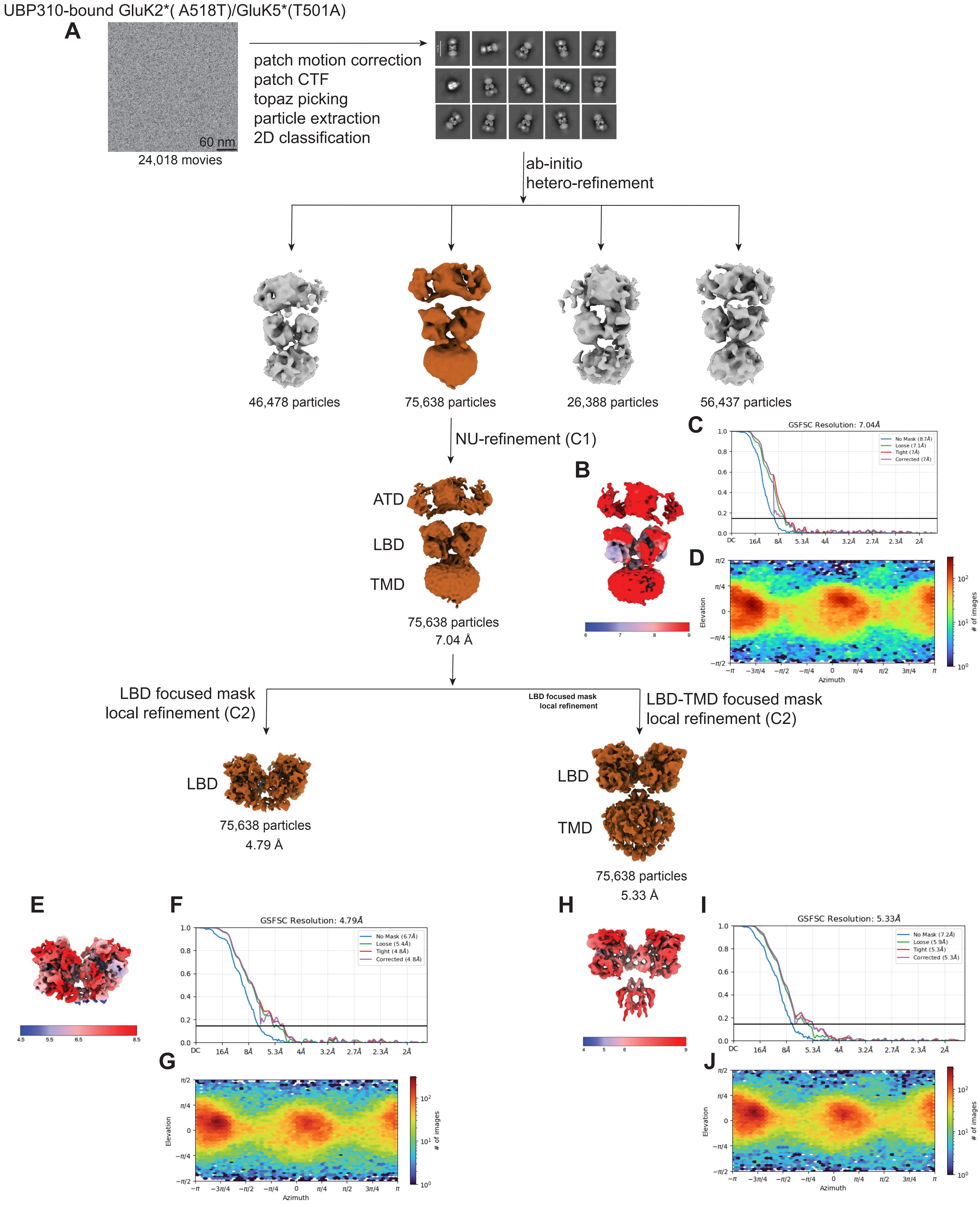
UBP310-bound GluK2*( A518T)/GluK5*(T501A)
A
60 nm
24,018 movies
patch motion correction
patch CTF
topaz picking
particle extraction
2D classification
ab-initio
hetero-refinement
46,478 particles
75,638 particles
26,388 particles
56,437 particles
NU-refinement (C1)
ATD
LBD
TMD
75,638 particles
7.04 Å
B
C
GSFSC Resolution: 7.04Å
D
LBD focused mask
local refinement (C2)
LBD focused mask
local refinement
LBD-TMD focused mask
local refinement (C2)
LBD
75,638 particles
4.79 Å
LBD
TMD
75,638 particles
5.33 Å
E
F
GSFSC Resolution: 4.79Å
G
H
I
GSFSC Resolution: 5.33Å
J
Elevation
Azimuth
### of images

**Figure S8. Cryo-EM data processing workflow for the UBP310-bound GluK2*(A518T)/GluK5*(T501A) KARs.** **(A)** CryoSPARC-based data processing flowchart. **(B, E, H)** Local resolution estimation of the cryo-EM maps for the respective structures. **(C, F, I)** FSC curves of half- maps for full-length, LBD and LBD-TMD, respectively. **(D, G, J)** Particle angular distribution for the final 3D reconstruction for the respective maps.

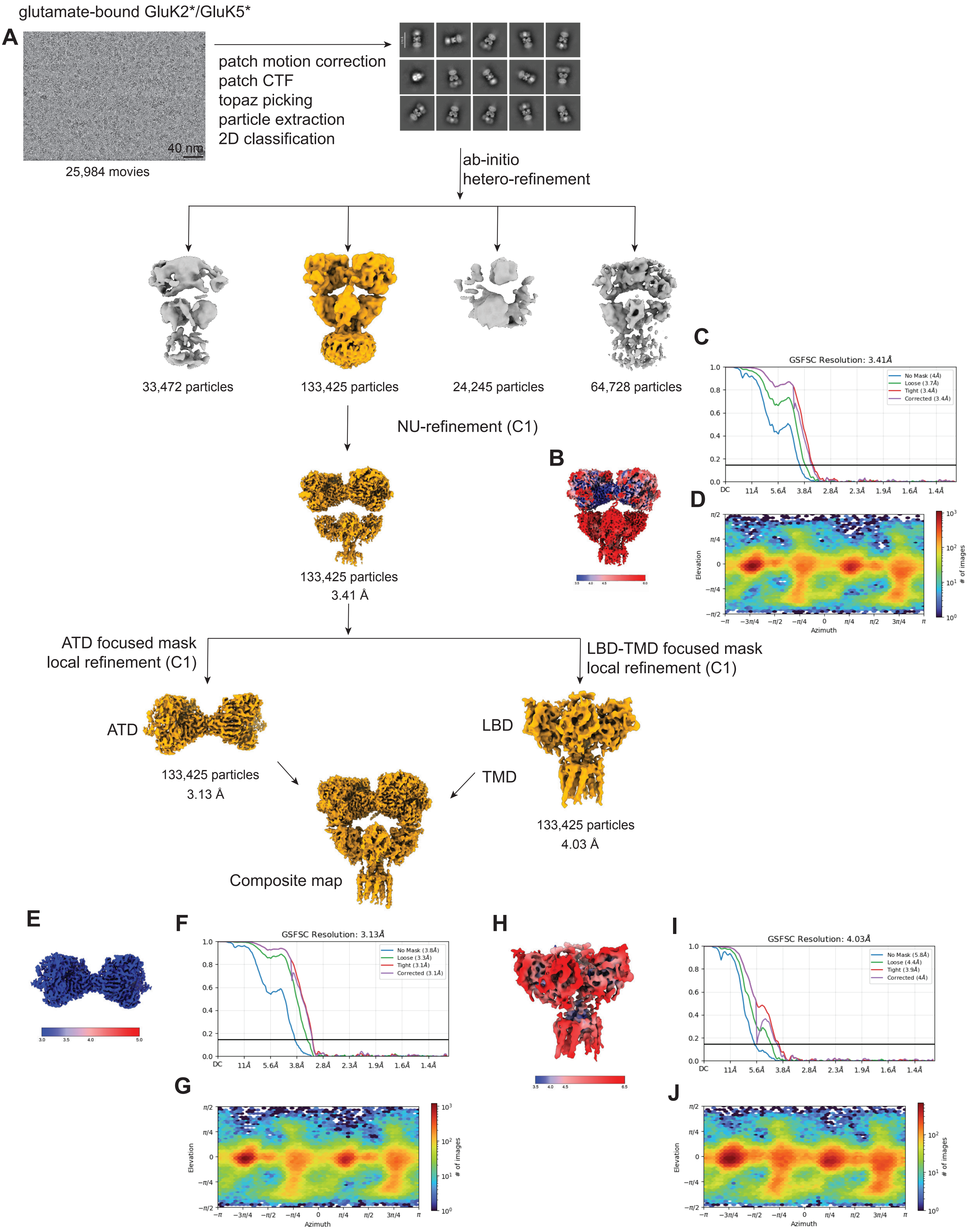
glutamate-bound GluK2*/GluK5*
A
40 nm
25,984 movies
patch motion correction
patch CTF
topaz picking
particle extraction
2D classification
ab-initio
hetero-refinement
33,472 particles
133,425 particles
24,245 particles
64,728 particles
NU-refinement (C1)
133,425 particles
3.41 Å
B
C
GSFSC Resolution: 3.41Å
No Mask (4Å)
Loose (3.7Å)
Tight (3.4Å)
Corrected (3.4Å)
D
Elevation
Azimuth
### of images
ATD focused mask
local refinement (C1)
LBD-TMD focused mask
local refinement (C1)
ATD
LBD
TMD
133,425 particles
3.13 Å
133,425 particles
4.03 Å
Composite map
E
F
GSFSC Resolution: 3.13Å
No Mask (3.8Å)
Loose (3.3Å)
Tight (3.1Å)
Corrected (3.1Å)
G
H
I
GSFSC Resolution: 4.03Å
No Mask (5.8Å)
Loose (4.4Å)
Tight (3.9Å)
Corrected (4Å)
J

**Figure S9. Subunit-specific conformational changes in glutamate-bound GluK2*/GluK5* KARs.**
**(A)** CryoSPARC-based data processing flowchart. **(B, E, H)** Local resolution estimation of the cryo-EM maps for the respective structures. **(C, F, I)** FSC curves of half- maps for full-length, LBD and LBD-TMD, respectively. **(D, G, J)** Particle angular distribution for the final 3D reconstruction for the respective maps.

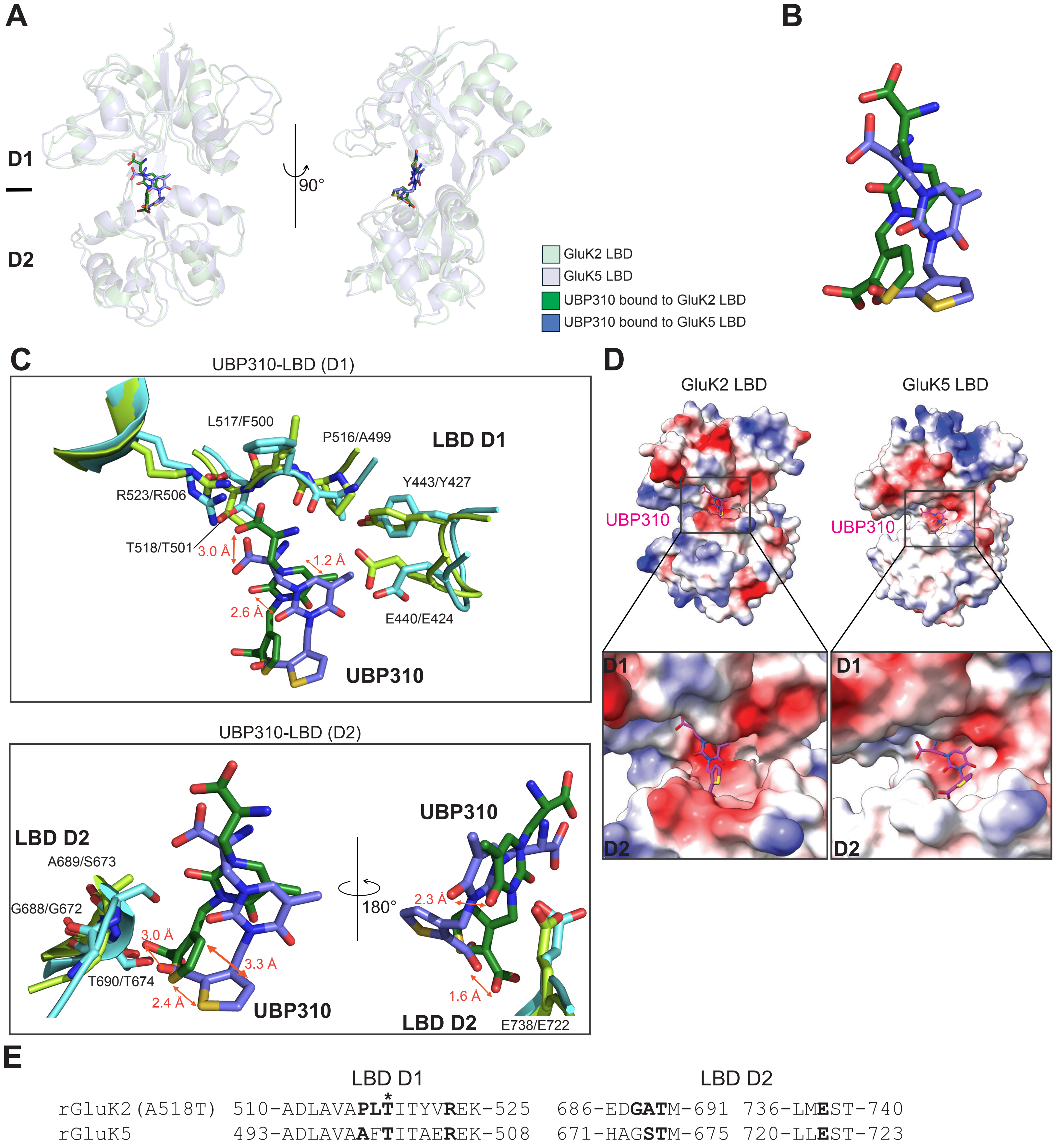
A
D1
D2
90°
GluK2 LBD
GluK5 LBD
UBP310 bound to GluK2 LBD
UBP310 bound to GluK5 LBD
B
C
UBP310-LBD (D1)
L517/F500
P516/A499
LBD D1
R523/R506
Y443/Y427
T518/T501
3.0 Å
1.2 Å
2.6 Å
E440/E424
UBP310
UBP310-LBD (D2)
LBD D2
A689/S673
G688/G672
3.0 Å
3.3 Å
T690/T674
2.4 Å
UBP310
180°
UBP310
2.3 Å
1.6 Å
LBD D2
E738/E722
D
GluK2 LBD
GluK5 LBD
UBP310
UBP310
D1
D2
D1
D2
E
LBD D1
LBD D2
rGluK2(A518T) 510-ADLAVAPLTITYVREK-525 686-EDGATM-691 736-LMEST-740
rGluK5 493-ADLAVAAFTITAEREK-508 671-HAGSTM-675 720-LLEST-723

**Figure S10. Structural comparison of UBP310 antagonist binding to the GluK2 and GluK5 LBDs.**
**(A)** Comparison of the GluK2 LBD (light green) and GluK5 LBD (light blue) subunits in complex with UBP310, aligned by their respective D1 lobes. The ligands are shown in stick representation (dark green for GluK2-bound, purple for GluK5-bound). **(B)** An overlay of UBP310 as bound to GluK2 (green) and GluK5 (purple). **(C)** Comparison of UBP310 binding in the D1 (top) and D2 (bottom) LBD lobes. Measured distances between UBP310 molecules in the ligand binding pockets (red) highlight the divergent orientations of UBP310 within the GluK2 versus GluK5 LBD binding pockets. The interacting residues within the GluK2 and GluK5 binding pockets are indicated (labeled as GluK2/GluK5). **(D)** Surface area of the GluK2 and GluK5 LBDs colored by electrostatic potential (red: -10kT/e; blue: +10 kT/e). Insets provide a magnified view of the ligand binding sites, showing the UBP310 molecule (magenta sticks) accommodated within the pocket. **(E)** Protein sequence alignment of the rat GluK2 (rGluK2) and rat GluK5 (rGluK5) LBDs. Residues involved in UBP310 binding are bolded. The asterisk (*) denotes the A518T mutation site in the rGluK2 construct used for cryo-EM.

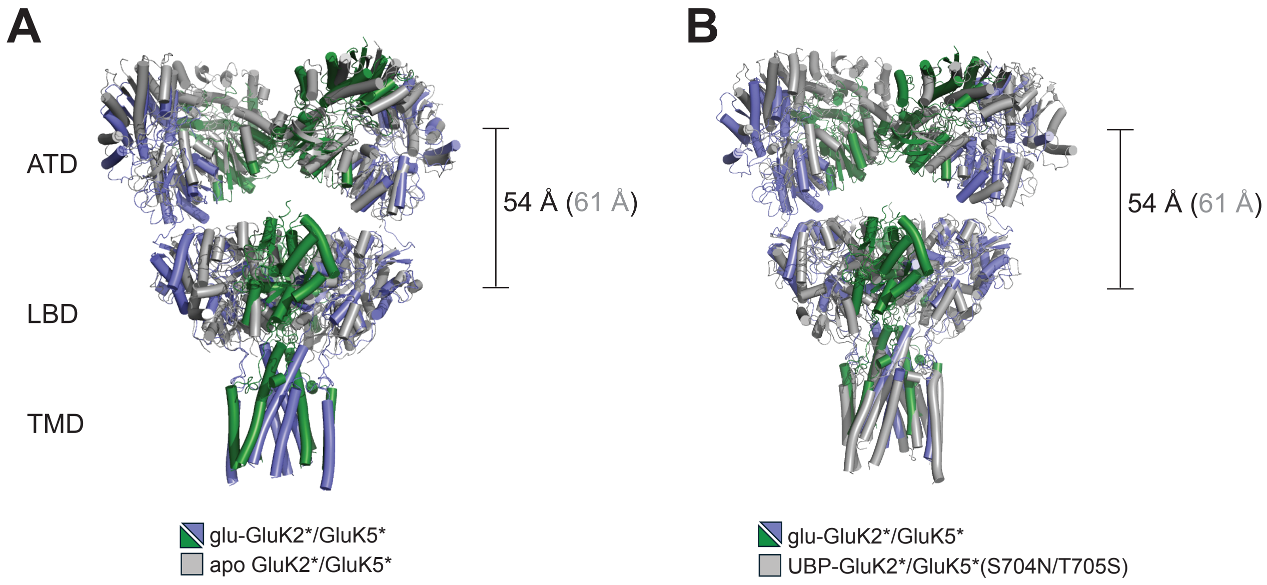


**Figure S11. Relative positioning of the ATD with respect to the LBD layer in the GluK2*/GluK5* KARs.**
**(A)** Superimposition of the apo and glutamate-bound GluK2*/GluK5* receptors. The distance bracket indicates the separation between the COM of the ATD and LBD layers in both states. **(B)** Superimposition of the glutamate-bound GluK2*/GluK5* and UBP310-bound GluK2*/GluK5*(S704N/T705S) receptors.

**Table S1. Cryo-EM data collection, refinement, and validation**

**(1) apo GluK2/GluK5 and glutamate-bound GluK2/GluK5**

| | Apo GluK2/GluK5 ATD-LBD<br><br>PDB: 9Z86<br>EMD-73887 | Apo GluK2/GluK5 LBD<br><br>PDB: 9Z85<br>EMD-73886 | Glutamate-bound GluK2/GluK5 ATD<br>PDB: 9Z8A<br>EMD-73891 | Glutamate-bound GluK2/GluK5 LBD-TMD<br>PDB: 9Z8B<br>EMD-73892 | Glutamate-bound GluK2/GluK5 full-length<br>PDB: 9C8Z<br>EMD-73893 | Glutamate-bound GluK2/GluK5 composite map<br>PDB: 9Z8D<br>EMD-73894 |
|---|---|---|---|---|---|---|
| **Data collection and professing** | | | | | | |
| Microscope | Titan Krios | Titan Krios | Titan Krios | Titan Krios | Titan Krios | Titan Krios |
| Microscope Camera | K3 | K3 | K3 | K3 | K3 | K3 |
| Energy filter | Gatan | Gatan | Gatan | Gatan | Gatan | Gatan |
| Magnification | 130,000 | 130,000 | 130,000 | 130,000 | 130,000 | 130,000 |
| Energy filter slit width (eV) | 20 | 20 | 20 | 20 | 20 | 20 |
| Collection software | SerialEM | SerialEM | SerialEM | SerialEM | SerialEM | SerialEM |
| Voltage (kV) | 300 | 300 | 300 | 300 | 300 | 300 |
| Electron exposure (e–/Å2) | 40 | 40 | 40 | 40 | 40 | 40 |
| Exposure rate (e–/Å2/frame) | 1 | 1 | 1 | 1 | 1 | 1 |
| Defocus range (μm) | -1.0 ~ -1.4 | -1.0 ~ -1.4 | -1.0 ~ -1.4 | -1.0 ~ -1.4 | -1.0 ~ -1.4 | -1.0 ~ -1.4 |
| Pixel size (Å) | 0.64 | 0.64 | 0.64 | 0.64 | 0.64 | 0.64 |
| Symmetry imposed | C1 | C2 | C2 | C1 | C1 | |
| Initial particle images (no.) | 135,217 | 135,217 | 535,217 | 535,217 | 535,217 | 535,217 |
| Final particle images (no.) | 56,821 | 56,821 | 133,524 | 133,524 | 133,524 | 133,524 |
| 0.143 FSC half map masked (Å) | 4.39 | 3.54 | 3.18 | 4.24 | 3.56 | |
| 0.143 FSC half map unmasked(Å) | 5.89 | 3.60 | 3.30 | 4.43 | 3.76 | |
| **Refinement** | | | | | | |
| Refinement package | Phenix | Phenix | Phenix | Phenix | Phenix | Phenix |
| Initial model used (PDB code) | 7KS0 | 7KS0 | 7KS3 | 7KS3 | 7KS3 | |
| 0.5 FSC model resolution masked (Å) | 6.71 | 3.75 | 3.32 | 4.40 | 6.41 | 3.86 |
| 0.5 FSC model resolution unmasked (Å) | 6.97 | 3.85 | 3.39 | 5.90 | 6.20 | 3.96 |
| Map sharpening B factor (Å2) | -91.1 | -85.9 | -103.6 | -170.3 | -106.3 | |
| Model composition | | | | | | |
| Non-hydrogen atoms | 20,611 | 8,203 | 12,168 | 11,758 | 21,111 | 24,000 |
| Protein residues | 2,567 | 1,023 | 1,544 | 1,473 | 2,653 | 3,027 |
| Ligands（NAG） | 14 | 0 | 0 | 6 | 0 | 5 |
| Ligands (UBA) | 0 | 0 | 0 | 0 | 0 | 0 |
| B factors ($A^2$) | | | | | | |
| Protein | 1052.60 | 153.72 | 175.08 | 312.4 | 50.38 | 38.18 |
| Ligand | 1052.60 | 0 | | 200.1 | | 37.59 |

| | | | | | | |
|---|---|---|---|---|---|---|
| R.m.s. deviations | | | | | | |
| Bond lengths (Å) | 0.003 | 0.003 | 0.003 | 0.003 | 0.003 | 0.003 |
| Bond angles (°) | 0.676 | 0.569 | 0.558 | 0.692 | 0.626 | 0.616 |
| Validation | | | | | | |
| MolProbity score | 2.41 | 2.38 | 2.69 | 2.61 | 2.50 | 2.63 |
| Clashscore | 14.92 | 11.65 | 11.27 | 13.56 | 12.25 | 14.42 |
| Rotamer outliers(%) | 3.16 | 5.61 | 5.98 | 5.49 | 4.09 | 5.15 |
| Ramachandran plot | | | | | | |
| Favored (%) | 94.93 | 96.52 | 90.76 | 93.82 | 93.28 | 93.58 |
| Allowed (%) | 5.07 | 5.48 | 9.24 | 6.18 | 6.72 | 6.39 |
| Disallowed (%) | 0 | 0 | 0 | 0.00 | 0.003 | 0.03 |
| CaBLAM outliers (%) | 2.62 | 1.31 | 4.52 | 3.05 | 3.44 | 3.75 |

#### (2) UBP310-bound $GluK2_{WT}/GluK5_{S704N/T705S}$ and $GluK2_{A518T}/GluK5_{T501A}$

| | UBP310-bound $GluK2_{WT}$/ $GluK5_{S704N/T705S}$ ATD PDB: 9Z87 EMD-73888 | UBP310-bound $GluK2_{WT}$/ $GluK5_{S704N/T705S}$ LBD-TMD PDB: 9Z88 EMD-73889 | UBP310-bound $GluK2_{WT}$/ $GluK5_{S704N/T705S}$ full-length PDB: 9Z89 EMD-73890 | UBP310-bound $GluK2_{A518T}$/ $GluK5_{T501A}$ LBD PDB: 9Z8E EMD-73895 | UBP310-bound $GluK2_{A518T}$/ $GluK5_{T501A}$ LBD-TMD PDB: 9Z8G EMD-73897 | UBP310-bound $GluK2_{A518T}$/ $GluK5_{T501A}$ full-length PDB: 9Z8H EMD-73898 |
|---|---|---|---|---|---|---|
| **Data collection and professing** | | | | | | |
| Microscope | Glacios | Glacios | Glacios | Glacios | Glacios | Glacios |
| Microscope camera | Falcon IV | Falcon IV | Falcon IV | Falcon IV | Falcon IV | Falcon IV |
| Magnification | 130,000 | 130,000 | 130,000 | 130,000 | 130,000 | 130,000 |
| Energy filter | Seletrix | Seletrix | Seletrix | Seletrix | Seletrix | Seletrix |
| Energy filter slit width (eV) | 10 | 10 | 10 | 10 | 10 | 10 |
| Collection software | EPU | EPU | EPU | EPU | EPU | EPU |
| Voltage kV) | 200 | 200 | 200 | 200 | 200 | 200 |
| Electron exposure (e–/Å2) | 40 | 40 | 40 | 40 | 40 | 40 |
| Exposure rate (e–/Å2/frame) | 1 | 1 | 1 | 1 | 1 | 1 |
| Defocus range (μm) | -1.0 ~ -1.4 | -1.0 ~ -1.4 | -1.0 ~ -1.4 | -1.0 ~ -1.4 | -1.0 ~ -1.4 | -1.0 ~ -1.4 |
| Pixel size (Å) | 0.91 | 0.91 | 0.91 | 0.91 | 0.91 | 0.91 |
| Symmetry imposed | C2 | C2 | C1 | C2 | C2 | C1 |
| Initial particle images (no.) | 655,658 | 655,658 | 655,658 | 415,968 | 415,968 | 415,968 |
| Final particle images (no.) | 95,364 | 153,265 | 153,265 | 75,638 | 75,638 | 75,638 |
| 0.143 FSC half map masked (Å) | 3.96 | 4.09 | 4.72 | 4.84 | 5.11 | 6.9 |
| 0.143 FSC half map unmasked(Å) | 4.07 | 4.14 | 4.93 | 4.94 | 5.29 | 6.8 |
| **Refinement** | | | | | | |
| Refinement package | Phenix | Phenix | Phenix | Phenix | Phenix | Phenix |
| Initial model used (PDB code) | 7KS0 | 7KS0 | 7KS0 | 7KS0 | 7KS0 | 7KS0 |
| 0.5 FSC model resolution masked (Å) | 4.08 | 4.15 | 4.92 | 5.16 | 5.74 | 7.09 |
| 0.5 FSC model resolution unmasked (Å) | 4.17 | 4.19 | 6.36 | 6.13 | 5.92 | 7.20 |
| Map sharpening B factor (Å2) | -133.4 | -147.4 | -208.7 | -253.5 | -286.3 | -414.6 |
| Model composition | | | | | | |
| Non-hydrogen atoms | 12,408 | 12,157 | 24,424 | 8,768 | 92,99 | 21,827 |
| Protein residues | 1,544 | 1,498 | 3,046 | 1,060 | 1,151 | 2,712 |
| Ligands（NAG） | 14 | 13 | 14 | 13 | 0 | 14 |
| Ligands (UBA) | 0 | 2 | 2 | 2 | 2 | 2 |
| B factors ($A^2$) | | | | | | |
| Protein | 193.09 | 208.67 | 308.6 | 416.91 | 359.12 | 1142.7 |
| Ligand | 191.73 | 226.32 | 307.04 | 221.21 | 154.54 | 1142.7 |
| R.m.s. deviations | | | | | | |
| Bond lengths (Å) | 0.003 | 0.004 | 0.002 | 0.004 | 0.003 | 0.003 |

| | | | | | | |
|---|---|---|---|---|---|---|
| Bond angles (°) | 0.643 | 0.638 | 0.609 | 0.799 | 0.712 | 0.70 |
| Validation | | | | | | |
| MolProbity score | 2.30 | 2.38 | 1.91 | 2.18 | 2.25 | 2.08 |
| Clashscore | 6.74 | 8.63 | 10.03 | 15.02 | 15.71 | 13.12 |
| Rotamer outliers(%) | 4.87 | 4.22 | 0.004 | 0.43 | 0.1 | 0.00 |
| Ramachandran plot | | | | | | |
| Favored (%) | 93.88 | 93.18 | 94.25 | 91.57 | 89.87 | 93.01 |
| Allowed (%) | 6.12 | 6.68 | 5.72 | 8.33 | 9.87 | 6.96 |
| Disallowed (%) | 0 | 0.13 | 0.03 | 0.1 | 0.26 | 0.04 |
| CaBLAM outliers (%) | 3.80 | 3.48 | 3.03 | 5.25 | 6.34 | 4.43 |
